# The Role of Cytonemes and Diffusive Transport in the Establishment of Morphogen Gradients

**DOI:** 10.1101/2024.12.22.629977

**Authors:** Jay Stotsky, Hans G. Othmer

## Abstract

Spatial distributions of morphogens provide positional information in developing systems, but how the distributions are established and maintained remains an open problem. Transport by diffusion has been the traditional mechanism, but recent experimental work has shown that cells can also communicate by filopodia-like structures called cytonemes that make direct cell-to-cell contacts. Here we investigate the roles each may play individually in a complex tissue and how they can jointly establish a reliable spatial distribution of a morphogen. To this end, we formulate models that capture fundamental aspects of various cytoneme-based transport mechanisms. In simple cases, exact solutions are attainable, and in more complex cases, we discuss results of numerical simulations.

## 1 Introduction and background

Pattern formation in embryonic development is a major, yet poorly understood, process in developmental biology. Throughout early development, maternal and zygotic cues regulate gene expression, cell proliferation, and differentiation in space and time in a highly-reproducible manner. Various theories have been proposed to explain how individual cells in an aggregate of essentially-identical cells differentiate into a collection of different cell types organized into the appropriate spatial pattern in response to factors called morphogens.

The most prominent of these are Turing’s theory (Turing 1952) and the theory of positional information. In Turing’s theory patterns arise from the interaction of reaction and transport by diffusion without external cues. This can explain certain patterns such as spots on insects, but is less useful when the spatial pattern is determined by a specified distribution of morphogens, and here the theory of positional information due to Child (1941) and Wolpert (1969) is more appropriate. In positional-information mechanisms of morphogen-mediated patterning, cells determine their spatial position by sensing the local levels of a spatially-graded morphogen, often produced at the boundary of a region to be patterned. The simplest model of how this functions is embodied in the French flag problem, which is to subdivide a finite interval [0, *L*] on the *x*-axis into three equal partitions, the first to be colored blue, the second white, and the third red. This is accomplished by producing a morphogen at the left boundary, the distribution of which then evolves via diffusive transport and first-order decay in the interior, and removal at the right boundary. This leads to a monotone-decreasing gradient of the morphogen level, and cells interpret the morphogen field via two thresholds that then lead to the appropriate coloration of the intervals. This mechanism has gained widespread appeal for interpreting the appearence of different cell types in an initially-unstructured tissue such as the *Drosophila* wing disc (Stapornwongkul & Vincent 2021), but it also raises numerous questions, such as how does the mechanism cope with tissues of different sizes (Umulis & Othmer 2013).

Both Turing’s theory and the theory of positional information rely on diffusion as the means of transport (Müller *et al*. 2013; Othmer & Scriven 1971; Stapornwongkul & Vincent 2021) but other means of communication in tissues have been discovered in recent years, and the one of interest here uses what are called cytonemes. Cytonemes are rod-like structures similar to filopods, which extend from a cell by polymerizing actin at the tip. They are 𝒪(0.1−0.5)*µm* in diameter and up to 100 *µm* in length. In the *Drosophila* wing disc they can be as long as 80 *µm*, which spans the entire length of the disc at early stages (Ramírez-Weber & Kornberg 1999). As with filopodia, cytonemes contain an actin network, but in addition, some contain myosin motors such as Myo-10 that can step along actin filaments to transport cargo (Hall *et al*. 2021). Connections between a cytoneme and a cell, or directly between cytonemes, leads to three forms of communication between cells, as shown in Figure 1.

**Figure 1:**
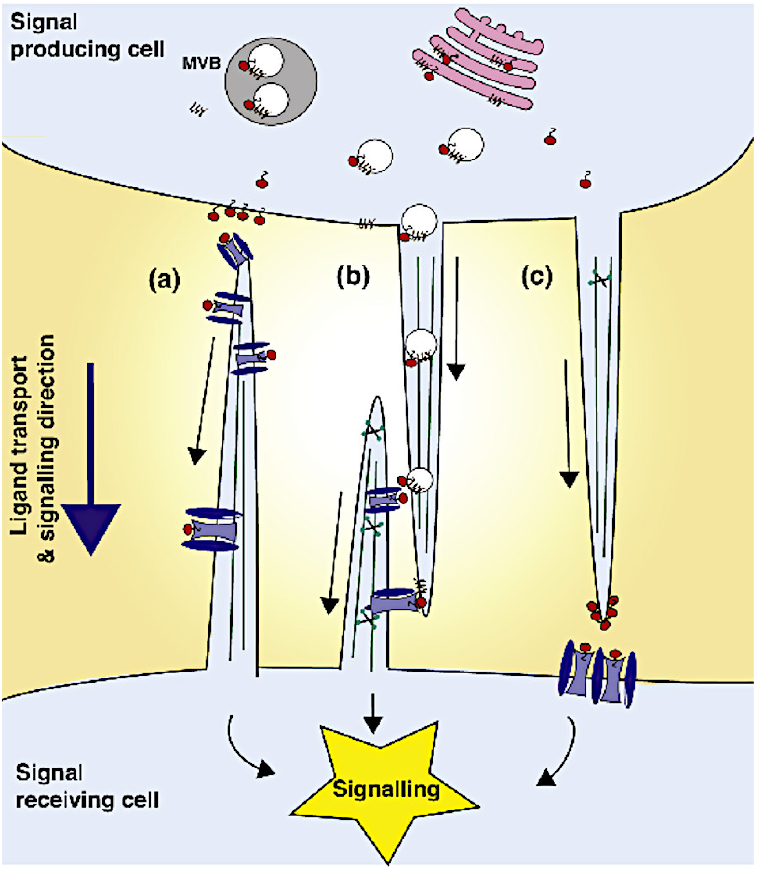
The three modes of cytoneme transport. (A) the RIT mode, (B) the ST mode, (C) the PIT mode. In general cells adopt only one of these modes. From (Zhang & Scholpp 2019)

In the first mode, labelled (a), the cytoneme is generated at a non-producing receiver cell such as in the wing disc of *Drosophila* (Stanganello & Scholpp 2016) and connects to a producer cell, from which it either receives a bolus of the morphogen Wg and then retracts, or receives multiple doses that are transported along the outside of the cytoneme. We call this receiver-initiated transport (**RIT**). In addition to transporting Wg, RIT is also used in the *Drosophila* wing disc to transport another important morphogen called Dpp (Zhang & Scholpp 2019). In a second type (b), both receiver and sender send out cytonemes that can create synapse-like connections to transfer morphogen from producer to receiver. We call this synaptic transport (**ST**), and this mode is used in the chick limb bud (Kornberg 2013). Finally, a third method (c), is used in vertebrates to transfer the morphogen Wnt from producers to receivers, and we call this producer-initiated transport (**PIT**). In this mode a morphogen-producing cell extends a cytoneme that attaches to a receiver cell and transfers a bolus of morphogen to the receiver. In some cases there may be successive transfers of morphogen-filled vesicle-like structures carried along the cytoneme by molecular motors.

One sees that PIT is more direct since the receiver gets the signal upon contact with the cytoneme, whereas in the RIT mode, if a bolus is transferred the receiver must extend and retract the cytoneme to obtain the signal. Why Wnt transport in vertebrates uses the former whereas Wg transport in invertebrates is via the latter remains an open question.

One of the first experimental studies of the role of cytonemes showed how air-sac primordia in the *Drosophila* wing disc extend cytonemes to both Fgf- and Dpp-secreting cells (Ramírez-Weber & Kornberg 1999). Recent work shows that they play a role in a wide variety of systems, including development of neuron types in the mouse neural tube (Hall *et al*. 2021) and in Hh transport in the wing disc (Bischoff *et al*. 2013). At present the available experimental information is relatively high-level, demonstrating the existence of cytonemes and what their cargo is, but details on what controls the origination and number of cytonemes per cell in either the PIT or RIT mode remain sparse. Do extracellular signals bias the search process, or is the process random? How do cytonemes remain in contact during either loading or unloading of their cargo, and when vesicles are involved in PIT cytonemes, what determines when the transfer process stops? A number of recent reviews cover the variety of systems in which cytonemes play a role (Daly *et al*. 2022; Korenkova *et al*. 2020; Casas-Tintó & Portela 2019; González-Méndez *et al*. 2019; Routledge & Scholpp 2019; Yamashita *et al*. 2018), but much remains to characterize the details of the processes involved.

Nonetheless, a number of models of communication and pattern formation based on cytonemes have been formulated. One of the first is that due to Vasilopoulos and Painter (VP) (2016), in which the authors develop and analyze a model for fixed direct contacts between signaling and receiving cells. Their focus was on the effect of long-range signaling in lateral-inhibition mechanisms of the type used in Notch-Delta signaling, and in their model the signaling network is static and a weight function is used to define who gives what and who gets what. They show that a variety of new pattern types can be generated, including sparse patterns of isolated cells and large clusters or stripes. Other models have been based on static or dynamic extension or retraction of cytonemes and the resulting search process. Models similar to the VP model were analyzed in Teimouri and Kolomeisky (2016), where *N* cytonemes were statically-attached to cells in a one-dimensional array, and a deterministic rate of transport was determined by the distance from the source cell. The authors showed how this could lead to a spatial gradient in the array. Stochastic transport of packets of morphogen were incorporated in Bressloff & Kim (2018) and Kim & Bressloff (2018), in which the authors described transport by a velocity jump process widely used to describe the movement of cells and organisms (Othmer *et al*. 1988; Hillen & Othmer 2000; Othmer & Hillen 2002; Hillen & Painter 2009). Reversal of the direction of packets was incorporated in the model, but since packets are carried along actin fibers by myosin motors (Hall *et al*. 2021), pauses are admissable, but reversals of direction may not be. More recently this model was applied at the level of a single cytoneme, which can pause and reverse, in a two-dimensional array of target cells (Bressloff & Kim 2019). A similar model was used to compare the steady-state gradient of morphogen resulting from cytoneme transport with that resulting from diffusion (Fancher & Mugler 2020), and recently computational models were used to analyze morphogen gradient establishment using cytonemes (Aguirre-Tamaral & Guerrero 2021; Rosenbauer *et al*. 2020).

While the details will be discussed later, we begin with a simple comparison between the diffusion-based transport and cytoneme-based transport models. To make this comparison, we first observe that for a single morphogen, the morphogen-producing cells often lie along a linear axis, and that the morphogen distribution does not vary strongly along this axis except near tissue boundaries. Thus, to begin, we reduce the problem to one-dimension. In this setting, each position *x* describes a column of tissue at a certain distance from the producing cells.

To make a fair comparison, we suppose that in each case, there is a specified mass flux *J* at *x* = 0. For diffusion this corresponds to the release rate of the morphogen, while for the cytoneme-based transport it corresponds to the rate at which cytonemes are generated, times the amount of morphogen each carries. Let *D* be the diffusion coefficient, *v* the cytoneme velocity, and *µ* a degradation coefficient. Then using the multi-cytoneme version of the PIT-cytoneme model discussed later one finds that for cytonemes

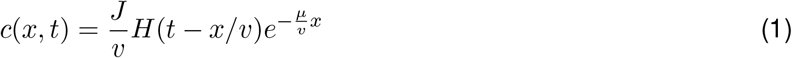

whereas for diffusion one finds that

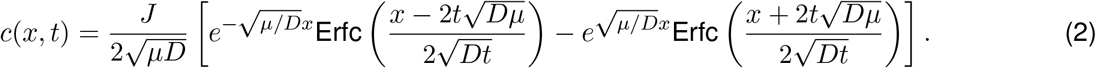

An important observation is that the long-time limits of both equations are exponential:

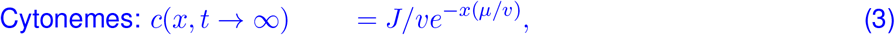

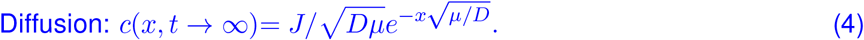

Thus, whenever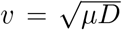, both solutions are equivalent in the limit *t* → ∞ suggesting that in biological situations where the diffusivity, advective velocity, and reaction rates are of roughly equal importance, either strategy could be applied in theory, if cells can achieve the same flux rate *J* for both transport mechanisms. On the other hand, the time-dependent dynamics are quite different. Unlike diffusion, the cytoneme-based transport generates a wave-like propagation pattern, but since both modes yield similar long-term behavior, either is viable for morphogen transport and may be used concurrently.

As we show herein, at a certain level of description cytoneme transport is simple and direct, even in complex tissues, whereas diffusive transport frequently involves intermediate steps such as binding to receptors or cell-to-cell transport. The latter form of signalling is called transcytosis or juxtracine signalling, and is clearly different from signalling via cytonemes. In Section 2, our first objective is to construct minimal models of the the PIT and RIT cytoneme-based transport **(CBT)** modes. Despite their simplicity, these models highlight key features of the transport, including spatial profiles of morphogen distribution, how morphogen is taken up by cells, and the presence of key parameter groups that govern the shape of the spatial distributions of morphogens. These models are also useful as precursors for more complicated models. After introducing the models, we discuss further applications of them, ranging from the single cytoneme setting to results for many cells and cytonemes.

The analysis in the first half of the paper provides a useful set of solvable problems and highlights some key issues that must be addressed in any cytoneme model, but they are limited in scope since nonlinearities are difficult to incorporate. In Section 4 we develop an algorithm for stochastic simulation of CBT and show that under appropriate conditions the analytical and numerical results agree. The stochastic simulation approach is very flexible and easy to extend, and we use it on a simple example involving biochemical feedback that cannot easily be treated analytically.

Several extensions to the models are discussed in Section 5. This includes more detailed modeling of puncta moving along a cytoneme, as well as how cell growth affects the distribution of morphogen that results from a CBT process. In Section 6 we compare diffusive-transport to CBT in a hexagonal array of cells that is used to represent an epithelial tissue such as the wing disc. Using the theory developed in (Stotsky *et al*. 2021), we show that the resulting transport is much more preciselydirected when cytonemes are used, leading to an advection-reaction equation along just one direction. In contrast, diffusion along the hexagonal edges of cells leads to a 2D diffusion under certain parameter scalings.

While there are many other possibilities not covered herein, our goal is to lay the groundwork for more detailed cytoneme modeling and simulation. As is elaborated below, more detailed modeling is currently challenging because there is little experimental evidence as to how cytonemes navigate their environment, how they attach to target cells, and how motor proteins shuttle cargo along cytonemes.

## 2 CBT via single cytonemes

### 2.1 An overview of the levels of description

A detailed mathematical model of the motion of a cytoneme should include components such as the biochemistry and mechanics involved in the creation and extension of a cytoneme, how they interact with the substrate and find the interstitial space to move between cells, how they decide which cells to attach to, and how morphogen is transported along a cytoneme and released at its terminus. At present there is insufficient biological information available to construct such detailed models, and we focus on simplified models of motion for which a high-level description can be constructed. These are sufficient to answer fundamental questions regarding the shape and time-scale of morphogen distributions arising from CBT, and predictions from these models could motivate future experimental studies. When further information on the details mentioned above becomes available, it can serve to relate parameters such as the cytoneme extension velocity in the simpler models to underlying biological processes such as myosin activation.

The motion of an individual cytoneme is fundamentally stochastic, but throughout we assume that cytonemes remain connected to a cell as they grow or shrink. Therefore the motion can be described by the motion of the tip, and in effect we treat cytonemes as point particles whose motion can be described by Newton’s laws. Of the three modes of cytoneme-based transport described earlier, we only consider PIT and RIT modes in this paper, and therefore assume that the tip of a cytoneme defines the time and location of attachments to distant cells. Analysis of the ST mode would require tracking the entire length of a cytoneme.

When there are stochastic variations in the cytoneme position and velocity this leads to the system of stochastic differential equations

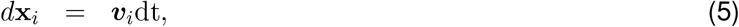

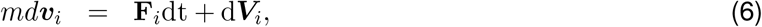

where (**x, *v***) ∈ *R*^*n*^, *n* = 1, 2, 3 are the positions and velocities (Othmer & Xue 2013). **F**_*i*_ represents the net imposed deterministic forces, *d***V**_*i*_ are stochastic forces. In the absence of stochastic forces, equations (5) & (6) are then the standard Newton equations for particles. In the presence of stochastic forces, these equations are stochastic differential equations. A commonly-used random process for modeling **V**_*i*_ is to assume that it is a Brownian motion, in which case *d***V** is Gaussian white noise. This can be used to model the random fluctuations in the choice of directions or speed of the tip, and under mild assumptions leads to a Fokker-Planck equation or a Smoluchowski equation (Othmer & Xue 2013).

Generation of new cytonemes is localized within cells and to describe this one could add a system of ordinary differential equations to (5) & (6) to describe the process in detail, but we simply impose some conditions on their appearance. We assume that new cytonemes are generated stochastically according to a Poisson process described later^1^, and thus the starting times (*t*_*s*_) for the motion of cytonemes are also random variables. Use of a Poisson process is justified when the start times between subsequent cytonemes are uncorrelated and when cells can support the creation of multiple cytonemes. In addition, cytonemes may enter into different states, e.g. motile and resting, and the transitions between these two states can also be described as a Poisson process. Experimental observation of the behavior of Myosin-X motors, which are important in cytoneme motion, indicates that this choice is realistic (Ropars *et al*. 2016).

While equations (5) and (6) apply to individual cytonemes, it is ultimately the combined effects of many cytonemes that results in graded morphogen distributions. This requires modeling the amount of morphogen transported by populations of cytonemes. In general, scaling up from an individual model to a population-level model is difficult^2^. However, when the generation times of new cytonemes are independent and exponentially-distributed, the conversion from individual cytoneme behavior to population-level behavior is straightforward and can be computed analytically as described in Appendix 8.2 and later on in the main text.

The amount of morphogen received by cell *i* at position *x* for *t* ∈ [0, *t*] can be described by a sum of the form

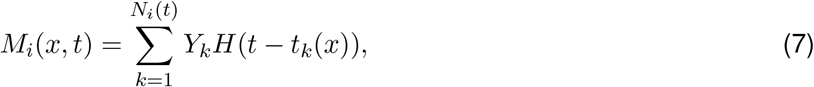

where the amplitudes *Y*_*k*_ are independent random variables representing the amount of morphogen in each cytoneme or each punctum on a cytoneme, *H* is the unit-step function, and *N*_*i*_(*t*) is a homogeneous Poisson counting process with parameter *λ* that counts the number of jumps in [0, *t*], assuming that *N* (0) = 0 with certainty. The arrival time *t*_*k*_(*x*) for PIT is the emergence time of the cytoneme plus the time needed to reach *x*. When multiple puncta are delivered to a given location *x* there are multiple *t*_*k*_ associated with *x*.

To develop models that are analytically solvable, we begin with movement in one space dimension at a constant macroscopic velocity, but later allow for pausing and other biologically-relevant processes.^3^ We further assume that the cells do not move, and thus the cytoneme’s tip position is governed by an equation of the form (5). As a result, the motion of a cytoneme can be described by the probability density function *p*(*x, t*) associated with (5) that represents the probability density that the tip of a cytoneme is in the interval (*x, x* + *dx*) at time *t*, and this must satisfy a boundary condition describing the emergence of cytonemes at *x* = *x*_0_. Later we also consider how things change when cells move with a velocity that is independent of how much morphogen they receive.

### 2.2 Producer-initiated transport on a single cytoneme

PIT is the most direct transport method, in that the cytoneme moves unidirectionally at a constant speed until it reaches a receiver cell, whereupon it stops and releases a morphogen payload which is internalized by the receiver cell. In PIT the transfer is thought to involve surface receptors, which means that there may be local loss of signals if the bolus saturates the available receptors.

As stated earlier, it is assumed that cytonemes are generated by a Poisson process with parameter *λ*, and therefore the inter-arrival times are exponentially-distributed. In this section the generation process is stopped after one cytoneme emerges, but later we consider what happens when this restriction is removed. We further assume that the process governing when cytonemes stop is also Poisson, and define *µ* to be the rate parameter for this process. It then follows that the density function *p*(*x, t*), so defined that *p*(*x, t*)*dxdt* is the probability of finding the tip in (*x, x* + *dx*) *×* (*t, t* + *dt*), satisfies the first-order hyperbolic equation and boundary condition

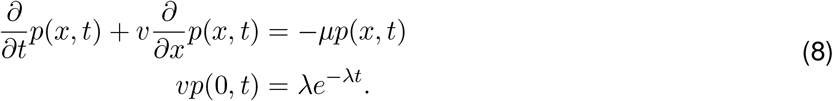

The solution of this is

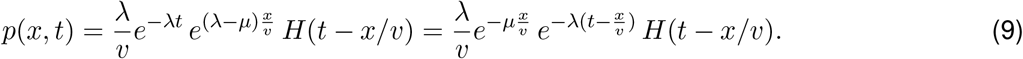

Each cytoneme starts at a random time 𝒯_*s*_ drawn from the distribution given by *p*(0, *t*), and eventually halts at a random time, 𝒯_*s*_ + 𝒯_*h*_ where 𝒯_*h*_ is exponentially-distributed with rate *µ*. Since the cytoneme simply advects forward at velocity *v* after it is generated, the distance traveled, *x*^*^ = *v*𝒯_*h*_ can be computed once 𝒯_*s*_ and 𝒯_*h*_ are known. Thus the time and spatial distribution of where the morphogen bolus is delivered is dependent on the generation and halting rates for a cytoneme, but these are governed by independent processes.

For a single PIT-cytoneme delivering a unit morphogen payload^4^ over a distributed array of cells, equation (7) becomes

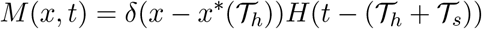

where *δ*(·) is the Dirac-delta function and *x*^*^(𝒯_*h*_) is the position of the cytoneme-tip at time 𝒯_*h*_; when the cytoneme moves at constant velocity *v*, this is *x*^*^ = *v*(𝒯_*h*_ − 𝒯_*s*_). The expectation of this quantity, denoted by *m*_1_(*x, t*), is the probability of a receiver-cell at position *x* having received a unit morphogen bolus by time *t*.^5^ With 𝒯_*s*_ and 𝒯_*h*_ drawn from exponential distributions of rates *λ* and *µ* respectively, this expectation can be found directly by computing

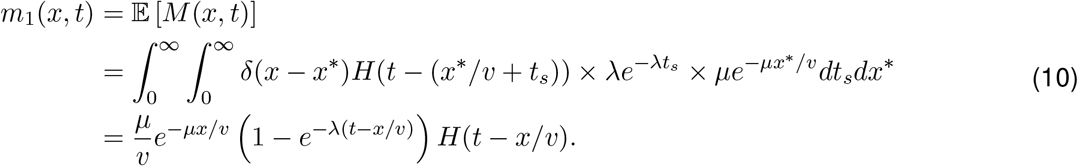

This solution is shown in Figure 3. As *t* → ∞ this approaches *m*_1_(*x*, ∞) = *µe*^−*µx/v*^, *i*.*e*., the expected position for delivery of the morphogen is exponentially-distributed. This coincides qualitatively with the distribution resulting under diffusion from a point with first-order decay.

The foregoing result can also be obtained by an alternative method, which makes direct use of equation (8), and later will prove useful for more complicated models. For a single cytoneme, the expectation *m*_1_(*x, t*) satisfies the recursion relation,

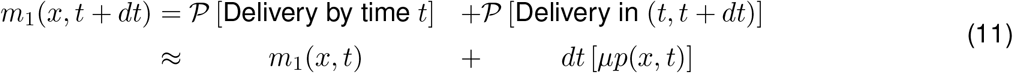

and therefore *m*_1_ satisfies

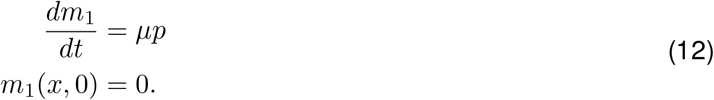

The solution is

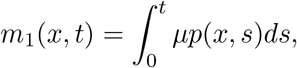

which is equivalent to the previous result for *m*_1_.

To summarize for the PIT model, let Λ(*t*) = *λe*^−*λt*^*H*(*t*) and 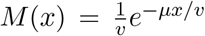 be the densities for the starting time and stopping points, respectively. Then it follows from equations (8) and (12) that

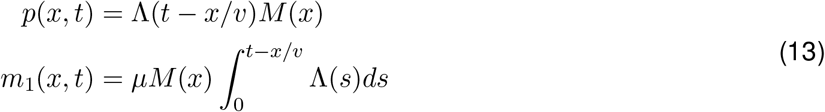

The ODE form for *m*_1_ is also useful since it is easy to futher include internal aspects of the cell behavior. For instance, if there is a linear rate of degradation of *m*_1_ inside a cell, then at any point in time, the amount of morphogen within a cell is governed by

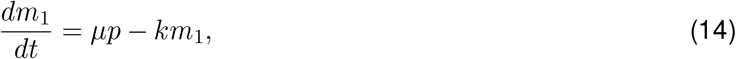

and therefore

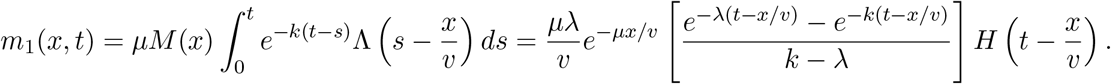

When internal dynamics alter the morphogen level, then *m*_1_ cannot be interpreted as the probability of the morphogen reaching a cell, but rather as the probability that the morphogen is currently in the cell, and has not yet been degraded. In the following we will usually compute only the accumulated morphogen (*i*.*e*., we set *k* = 0), but as shown above it is straightforward to extend the results as needed.

While the model described in equation (8), even when generalized as above, is very simple, it is already clear that an important factor in the morphogen distribution is the length scale *ℓ*^*^ = *v/µ*. This length essentially governs how quickly the cytoneme halts compared with its extensional velocity. Let *ℓ*_*c*_ denote the length of a single cell – then, for *ℓ*^*^ ≈ *ℓ*_*c*_ the distribution will be concentrated about *x* = 0, whereas for *ℓ*^*^ ≫ *ℓ*_*c*_ it will have a shallow gradient. Further, the value of *x*_1*/*2_ = ln 2*/ℓ*^*^ is the number of cell-lengths over which the probability of delivering morphogen is halved.

### 2.3 Receiver-initiated transport

In the RIT mode in which a single bolus is received, receiver-cells search for producers to extract morphogen, and the cargo in the cytoneme has direct access to the cytosol of the receiver cell.^6^ When the bolus is a morphogen, it can activate downstream signaling directly. In this RIT mode transport involves three main stages: i) the receiver-cell generates a cytoneme which searches for a producer cell, ii) the cytoneme attaches to the producer-cell, and iii) after loading it retracts to its orgin at a receiver-cell. Because there are multiple stages, simple RIT models are more complicated than the PIT models, which essentially involved one stage (two if morphogen dynamics along attached cytonemes are involved). A diagram of these stages is given in Figure 2.

**Figure 2:**
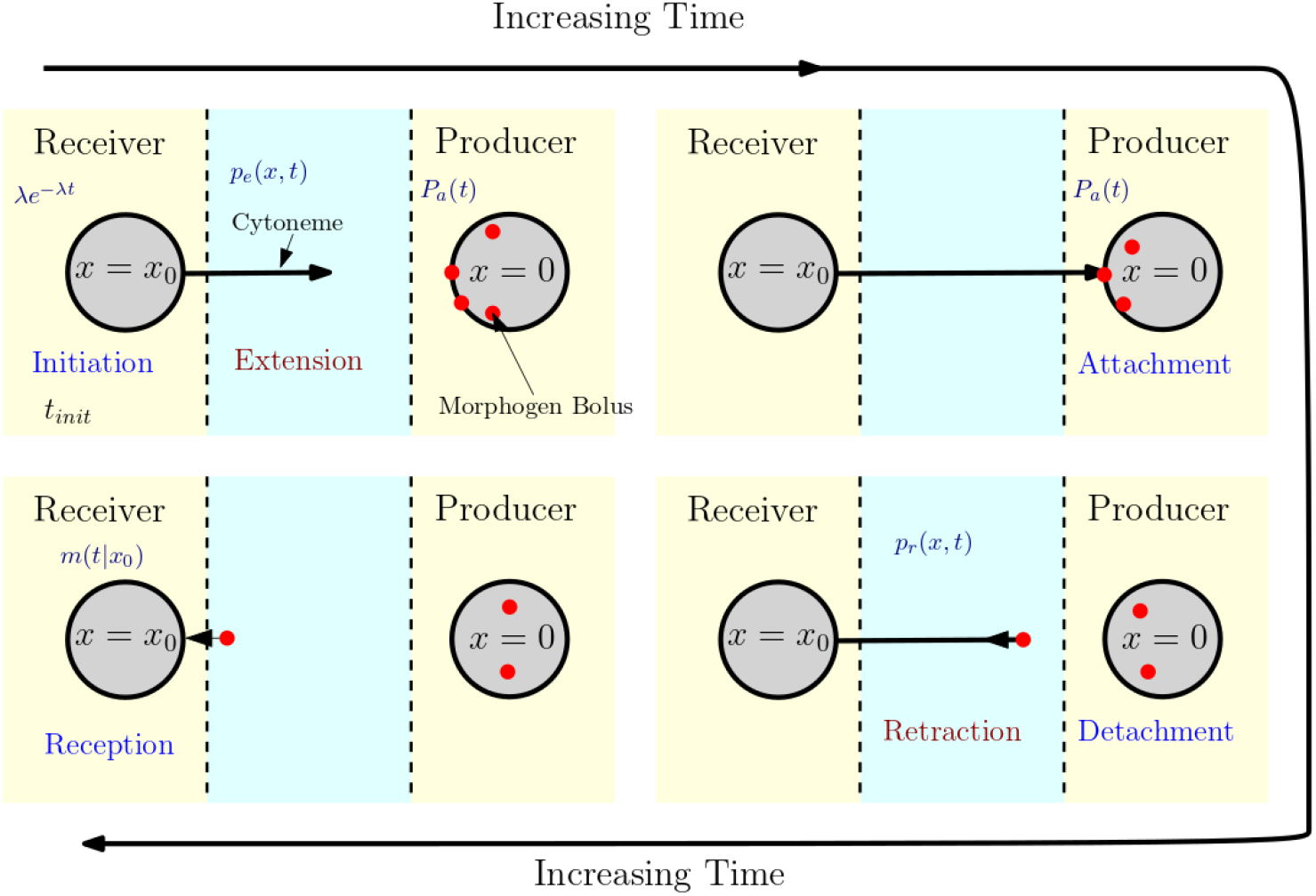
A diagram of the stages in the RIT mode of transport and the trajectory of the cytoneme tip. Cytonemes are first initiated by a receiver cell at position *x*_0_ (upper left), then must travel a distance to reach a producer cell at *x* = 0. They then attach, receive morphogen (upper right). Next, they begin retracting (lower right), until returning to their host cell where they deliver morphogen (lower left). Yellow backgrounds indicate steps involving stochastic waiting-times, and blue backgrounds with red stage names indicate transport steps. In the model below we assume that delivery occurs instantaneously upon return of the cytoneme.

To begin, we fix the receiver cell at *x*_0_ *>* 0 and assume that the producer cells are located at *x* = 0. We first consider a single cytoneme, generated at a random time, and in the first stage the cytoneme extends toward the producer. We assume that it reaches *x* = 0 with certainty and attaches instantaneously. Then the probability-density for the cytoneme to be located near *x* at time *t* on the inward trek is governed by

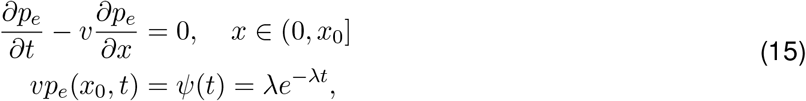

where *ψ*(*t*) is the probability density governing the cytoneme generation. The solution of this system is

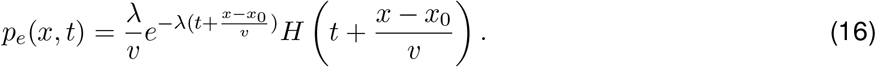

and the probability per unit of time for the cytoneme to reach the producer cell at *x* = 0 is *J*(*t*) = *vp*_*e*_(0, *t*). We further assume that while attached to a producer cell it spends an exponentially-distributed time with parameter *µ*_*d*_ loading, and thus the probability of attachment *P*_*a*_, evolves according to

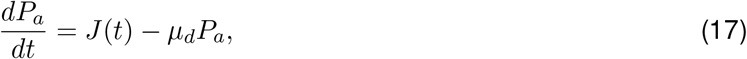

where *µ*_*d*_ is the detachment rate. This leads to the following expression for the probability that the cytoneme is attached at time *t*,

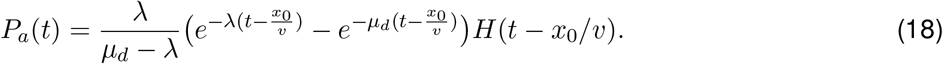

This is the density function for the two-step process of attaching and loading, wherein the time for each step is exponentially-distributed.

After detachment, the cytoneme is presumed to be carrying a morphogen packet^7^ and the probability density of the retreating cytoneme tip, *p*_*r*_ is governed by

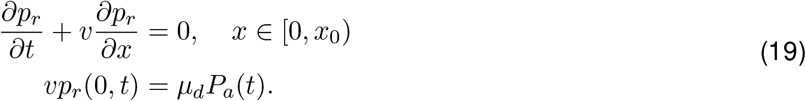

By combining these components we obtain the system

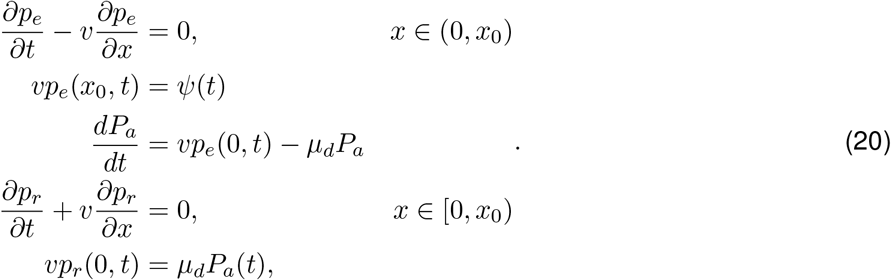

and from this we find that the expected morphogen accumulation in a cell at (*x*_0_, *t*) is

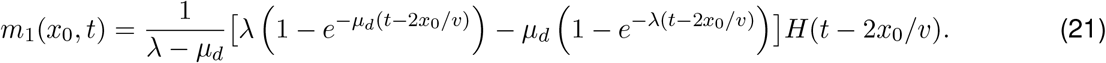

A plot of the RIT solution for *m*_1_ is given in Figure 3.

**Figure 3:**
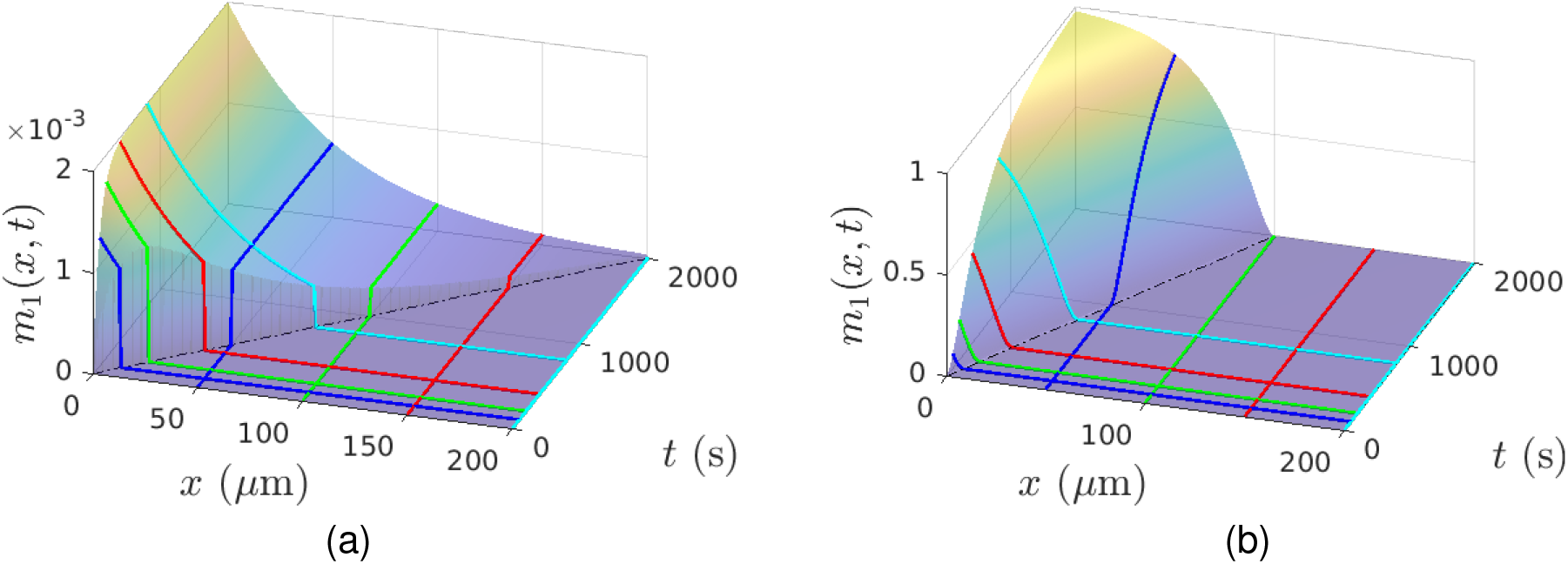
The morphogen accumulation for (a) PIT and (b) RIT models. Lines of constant *t* are highlighted to show the spatial distribution at *t* = 100 (blue), 200 (green), 400 (red), 800 (cyan) seconds. Since the solution of the RIT model only gives *m*_1_(*x, t*) for a specific *x*_0_, the spatial distribution shown is the result of a series of cells at all values of *x*_0_ simultaneously turning cytoneme generation on.

In contrast to the PIT process, the RIT process involves a start-up time of 2*x*_0_*/v* rather than *x*_0_*/v*. With all else equal, this leads to slower dynamics for the RIT process. On the other hand, since cytonemes are generated by the receiver cells, when the failure for the cytoneme to find a producer is small, this leads to greater morphogen accumulation over time at each receiver cell since each has its own *dedicated* cytoneme that will not halt at a different receiver cell.

The RIT solution can be further understood from the fact that the distribution of the sum of two exponentially distributed variables 𝒯_1_ + 𝒯_2_ with rates *α* and *β* is governed by the hypoexponential distribution, whose cumulative distribution function is

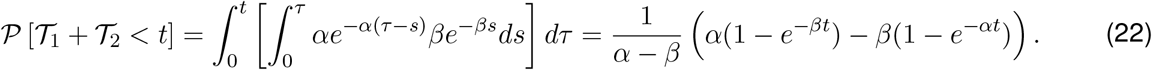

With *α* and *β* replaced by *λ* and *µ*_*d*_, and *t* shifted by the transit time, 2*x*_0_*/v*, we obtain *m*_1_ as before. Thus, the result is simply the sum of two exponential random variables shifted in time by the time interval for the cytoneme to extend and retract. This formula holds whether *α > β*, or *β > α*, and in the limit *α* = *β*, L’Hopital’s rule can be used to find the result,

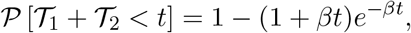

The whole solution has thus far been conditioned on the fact that the receiver-cell is a distance *x*_0_ from the producer cell. However, since *x*_0_ merely appears as a parameter throughout the model, the full solution can be obtained for any *x*_0_ *>* 0. Furthermore, if the cytonemes have a probability per unit time of failing during the extension phase, and turning back without morphogen, this modifies the equation for *p*_*e*_ as

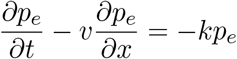

and the solution for the morphogen accumulation is simply multiplied by a factor 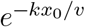 accounting for the failure probability, i.e.

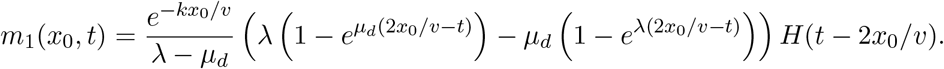

It is also straightforward to allow for the rate parameter of the cytoneme generation time to depend on *x*_0_, since this just implies that *λ* = *λ*(*x*_0_) everywhere that *λ* appears in the solution.

### 2.4 The effect of rest times in the RIT model

In reality, cytonemes do not simply extend for indefinite lengths of time at a particular velocity. Rather, they undergo intermittent stopping and starting, and the time spent in each resting phase has been suggested to be exponentially-distributed (Ricca & Rock 2010). To account for intermittent resting, we introduce a two-component model with *p*_*e*_(*x, t*) and *r*_*e*_(*x, t*) that represent cytonemes extending from a receiver-cell in the motile or rest phase. Since the transitions between these two states are exponentially-distributed, the RIT model above can be easily modified to allow for motile phases of mean duration *λ*_*m*_ and rest-phases of mean duration *λ*_*r*_. The equations are

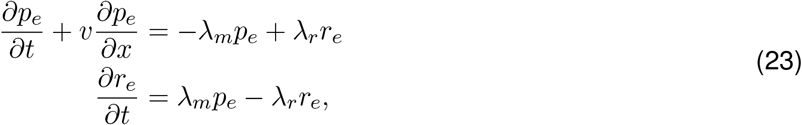

and if rest-phases can occur during extension as well as retraction, then we introduce a retraction rest phase, *r*_*r*_(*x, t*) and obtain equations for *p*_*r*_ and *r*_*r*_ similar to equation (23).

To account for cytonemes that fail to reach the producer cells, we assume that the time to failure is also exponentially-distributed, and thus modify the right-hand side of (23) with a term of the form −*k*_*d*_*p*_*e*_ to reflect this effect. By incorporating these assumptions one arrives at the system

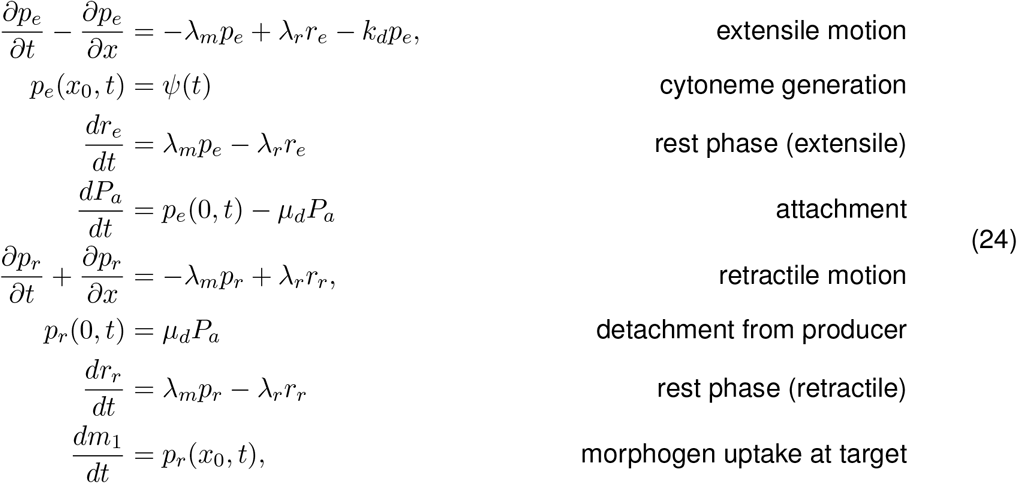

where again, the domain is (0, *x*_0_). Cytonemes that reverse prior to reaching the source region are not included in *p*_*r*_ since they do not contribute to the morphogen flux, but rather are simply eliminated via the degradation term *k*_*d*_*p*_*e*_. When *λ*_*m*_ = *λ*_*r*_ = 0, this model reverts to the standard RIT model defined in the previous section.

Upon non-dimensionalization one finds that the resulting system has four distinct groupings corresponding to *µ*_*d*_ as earlier, the time scales for entering and leaving the rest phase, as well as a time-scale associated with degradation. The resulting system can be solved via Laplace transform, but the inversion integral is complicated. Thus, we have solved the above system numerically using an upwind finite difference approximation to obtain approximate results for *m*(*x, t*) and the other quantities of interest.

The time dynamics of *m*_1_(*x, t*) are shown in Figure 4. When *λ*_*r*_ and *λ*_*m*_ are increased in tandem, the efficiency of the transport process is diminished by a factor that depends on the ratio *λ*_*m*_*/λ*_*r*_, which is the ratio of the mean time spent resting to the mean time spent moving. As these times increase, there is increased dispersion in the amount of time it takes for the cytonemes to travel and return due to the added source of stochasticity. When Λ = *λ*_*r*_ = *λ*_*m*_, as in Figure 4, transport is very fast when Λ = 0 (no resting phase) but slows significantly for Λ small, but then increases again for larger Λ. This can be understood as a result of two factors. First, for Λ ≈ 0, there is very little chance that the cytonemes enter a rest phase, so the transport is quite fast, essentially the same as if no rest phase were present. For Λ *>* 0 and moderately small, cytonemes tend to enter into the rest phase a small number of times, but then are trapped for a long time in each case because the exit rate *λ*_*m*_ = Λ is still low. For large Λ there are many exits and entrances from the rest state, and the cytonemes spend roughly half of their time resting, which leads to a roughly doubling of the time *t* needed to return to *x*_0_.

**Figure 4:**
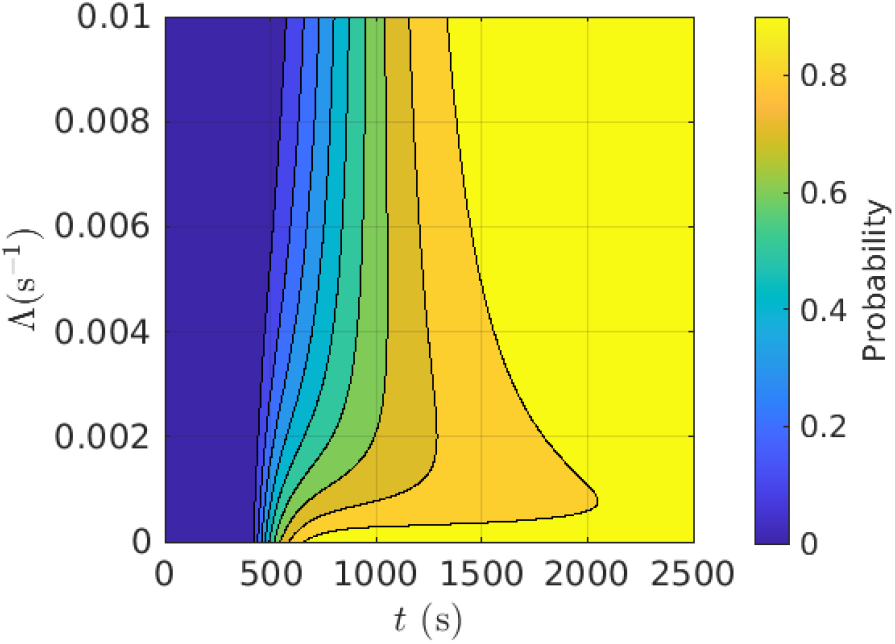
*m*_1_(20, *t*) as a function of Λ ≡ *λ*_*r*_ = *λ*_*m*_ and the return time to *x*_*o*_. For Λ = 0 the return time is 425 and for Λ *>* 0 and small it increases dramatically due to being trapped in long rest phases. For larger Λ, the velocity is effectively halved as many frequent transitions to and from resting occur.

Since the resting phase decreases the efficiency of cytoneme transport, a natural question is why cytonemes exhibit a resting phase, since it only seems to hinder the morphogen delivery process. While it is difficult to answer this conclusively, it may be that if cytonemes were not already highly optimized for efficiency, the resting phases would be much longer than those observed. Another aspect is that since the results are for one spatial dimension, the problem of orienting the cytoneme does not arise, whereas it does in a 2D or 3D tissue. It may be that during the resting phases the cytoneme is integrating various guiding queues to ensure that it travels towards its target. Extending the model to include various aspects of cytoneme transport is an area of future interest, though we do make some initial inroads into the importance of cytoneme transport direction in the stochastic model described in Section 4.

## 3 The effects of multiple cytonemes and packets per cytoneme

### 3.1 PIT and RIT with multiple cytonemes

The next step is to extend the previous models to multiple cytonemes, and multiple puncta on a cytoneme. A cautious reader might at first object to an extension to multiple cytonemes, since it appears that there could be interactions between them, which cannot occur with a single cytoneme, and thus is fundamentally different from the previous models. However, cytonemes are very thin, and recalling that the actual biological systems are 3D, we can assume that cytonemes can easily slip past each other by slight changes in their *y*− or *z*-coordinates without needing any modification to the model. This assumption could fail at very high cytoneme-densities, but as Figure 5 illustrates, this does not seem to be a problem in reality.

**Figure 5:**
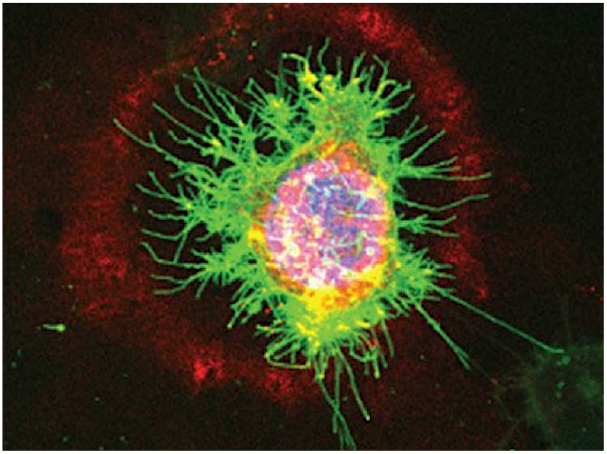
An image of a cluster of cytonemes that transport Wnt protein (red) between gastric cancer cells. The diameter of the red halo around these cells correlates with the average length of their cytonemes (green).Despite there being many cytonemes, it appears that it is uncommon for cytonemes to ‘collide’ with each other, and thus as described in the text, we assume that interactions are not a determining factor even when multiple cytonemes are present.Taken from (Mattes *et al*. 2018) with permission.

When multiple cytonemes are present the total amount delivered can be found by summing over single-cytoneme results conditioned on the sequence of times at which cytonemes are generated. Thus, instead of stopping the generation process after one cytoneme is generated, as in previous sections, suppose that new cytonemes are generated at random times {𝒯_*i*_} according to the distribution *ψ*(*t*) = *λe*^−*λt*^. This defines a Poisson process with constant rate parameter, *λ* (see Appendix 8.1 for a precise explanation), and implies that *N* (*t*), the expected number of cytonemes generated in [0, *t*), is a Poisson random variable with mean (*λt*)^−1^. The amount of morphogen received by each cell now depends upon the stochastic process governing the behavior of individual cytonemes, as well as the Poisson process that generates new cytonemes. As a result, the sum in (7) defines a compound Poisson process.

Since the distribution for an exponentially-distributed process resets at each occurrence, the inter-arrival interval times 𝒯_*i*+1_ − 𝒯_*i*_ for a Poisson process are independent random variables. Thus the morphogen delivery in a PIT process for a single cytoneme, which is given by

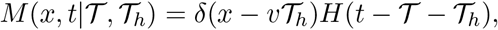

where 𝒯 and 𝒯 + 𝒯_*h*_ are the starting time and stopping times of the cytoneme, can be used to compute the sum in (7) as follows. In the following equations, *m*(*x, t*) is the *expected* amount of morphogen received per unit length of tissue from many cytonemes, rather than *m*_1_(*x, t*), which is the expected amount delivered at *x* by time *t* by a single cytoneme.

As is briefly outlined in Appendix 8.2, *m*(*x, t*) for PIT can be computed via a two-step conditioning process as

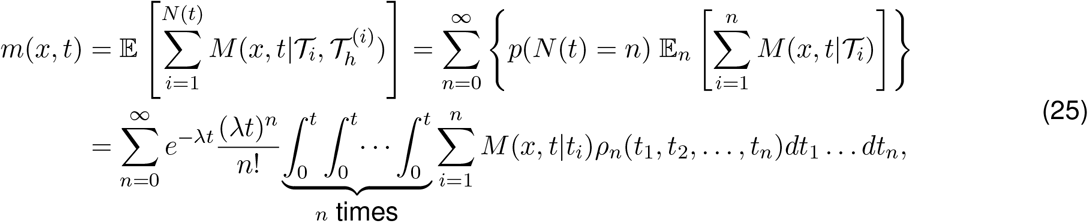

Where

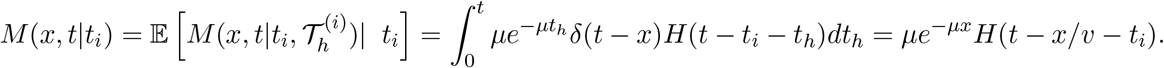

It follows from the theory of Poisson point processes that *ρ*_*n*_(*t*_1_, …, *t*_*n*_) is the density for events occurring at 𝒯_1_ ∈ (*t*_1_, *t*_1_ + *dt*_1_), etc., given that there are *n* events in [0, *t*). For a constant-rate process this reduces to

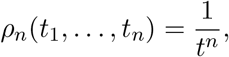

and therefore we can further simplify (25) by using the fact that *M* (*t, x*|*t*_*i*_) depends on only one of the *t*_*i*_’s. In particular,

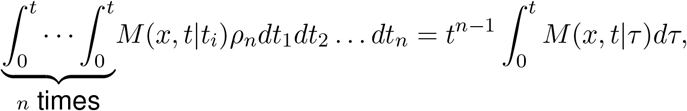

and this leads to

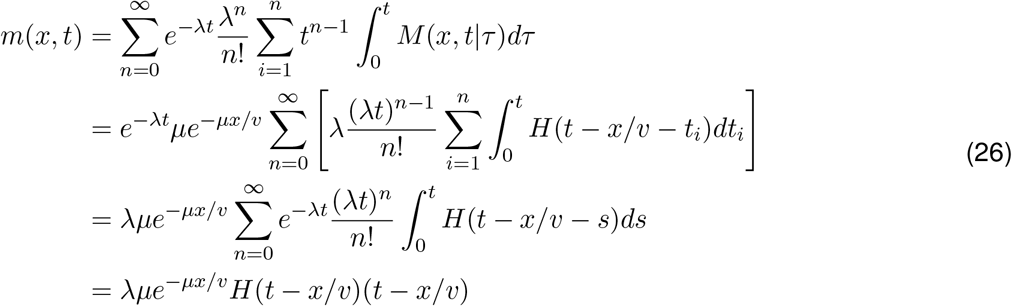

where the last step is possible since the integral is independent of *n*, and the remaining terms are simply a Taylor expansion for *e*^*λt*^.

While superficially similar to the previous result for *m*_1_(*x, t*), this result differs significantly in that *m*(*x, t*) grows indefinitely as *t* increases because we now have a continual series of cytonemes being extended (with generation times governed by a Poisson process) rather than just a single cytoneme. Nonetheless, comparing with earlier results shows that the mean amount of morphogen received at each position *x* is ultimately found directly by solving the original PIT and RIT model equations with *ψ*(*t*) replaced by *λH*(*t*). This is a direct result of the independence of the cytonemes, which implies that the governing equation for the cytoneme distribution is linear in *p*(*x, t*). The output of the PIT model under these conditions is shown in Figure 6.

**Figure 6:**
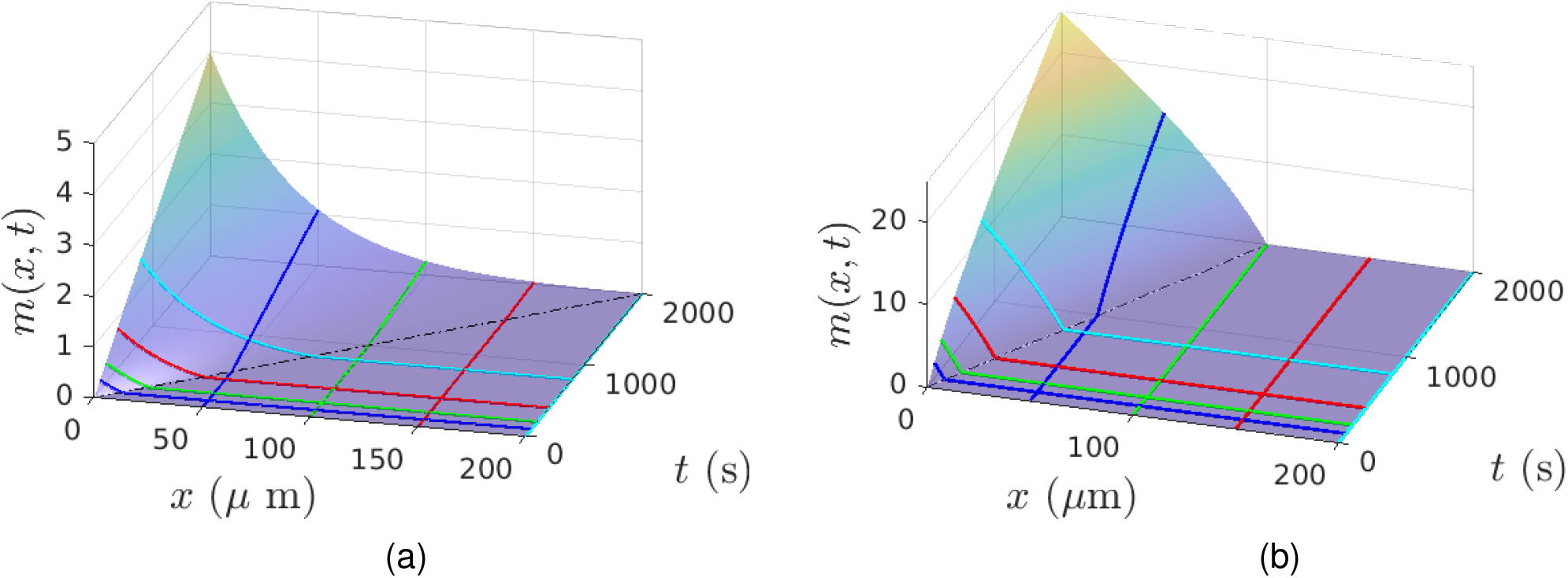
Expected morphogen accumulation *m*(*x, t*) for PIT **(a)** and RIT **(b)** with multiple cytonemes generated from Poisson process with rate *λ* = 0.01*/s* and *µ/v* = 100*µm*. Black dashed lines indicate the line *t* = *x/v* (left) and *t* = 2*x/v* (right) that govern the *x*-dependent time-delay prior to the beginning of accumulation in each case.

Similar features – a warm-up period of no accumulation, followed by a linear increase that has an *x*-dependent rate – is also present in the other forms of cytoneme-based transport. When *m* has a decay-term, as in equation (14), the result becomes

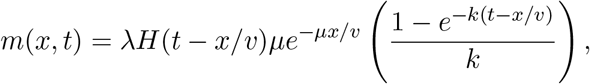

which reduces to the previous result when *k* → 0. In contrast to the *k* = 0 limit shown in Figure 6, for large times, *m*(*x, t*) remains bounded for non-zero *k*, in particular,

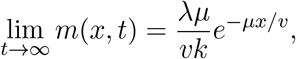

where the coefficient *λµ/vk* ties together cytoneme production, cytoneme halting, and degradation of morphogen inside the cells. Nonetheless, the spatial variation is still set by the length scale, *µ/v* independent of *λ* or *k*, which merely rescale the amount. This reflects the fact that transport of the morphogen is independent of processes inside the cells in this simple model. This also provides some indication that cell-level regulation may be more important in setting the morphogen-accumulation level, but not the shape of the resulting distribution. However, this result does not always hold if the cell behavior is more complicated, or involves a nonlinear response. For instance, when the cell uptake rate saturates at high densities of cytonemes, due for example to limited recepter availability, then we cannot assume that the rate of increase of *m* is proportional to *p*. For instance, if the uptake of morphogen through the cell membrane exhibits Michaelis-Menton-like binding kinetics, then

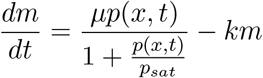

where *p*_*sat*_ is a saturation density, and the resulting solution is

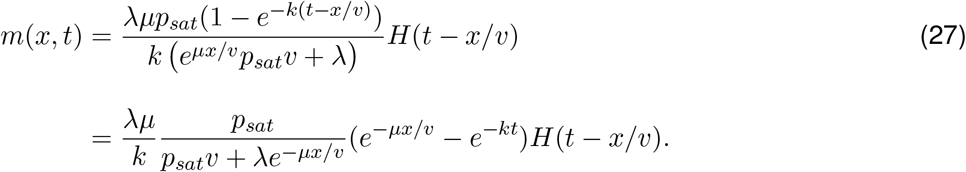

The long-time evolution is

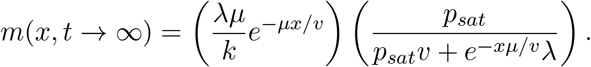

where the first factor is as before, and the second factor accounts for saturation dynamics. In these results the limit of no saturation occurs when either the saturation density *p*_*sat*_ becomes very large or *x* becomes large.

In contrast to diffusive transport mechanisms, the PIT model – whether for a single cytoneme or many – leads to a time-delay after startup during which *m*(*x, t*) is zero. For each *x*, once *t > x/v* the concentration increases exponentially in time with a coefficient dependent on that *x*, before eventually leveling off. To be precise, the diffusion equation is itself only an approximation of a process with finite velocity. However, nonetheless, the given the much smaller size of a single morphogen molecule to a cytoneme, we expect that the “first” morphogen molecule to arrive at its destination will occur before the first cytoneme arrives. However, the accumulation of morphogen may still be much slower since the vast majority of diffusing molecules will take extremely tortuous paths and have to bind/unbind many times prior to reaching the desired destination. In contrast, the molecules in a cytoneme have a much more simple route along a quasi-one-dimensional cytoneme.

The same approach can also be applied to the RIT model in equation (20). Under the condition that a single receiver cell located at *x*_0_ produces cytonemes at rate *λ*_0_ beginning at *t* = 0, we have *λ*(*x*_0_, *t*) = *λ*_0_*H*(*t*) where *H*(*t*) is the Heaviside step function, and the solution of equation (20) with this boundary condition is

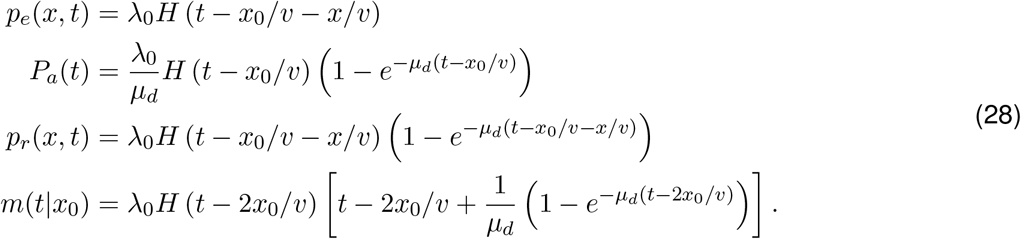

An example of the solution for this case is shown in Figure 6. Due to the finite velocity, there is an initial interval (0, 2*x*_0_) during which no morphogen is received, followed by gradual morphogen reception that depends on the attachment waiting-time distribution, and this gradually ramps up to a linear rate of increase with slope *λ*_0_. The linear increase rate is independent of *x*_0_, but the time-delay required to reach that rate is *x*_0_-dependent.

The results discussed thus far are applicable to single cytonemes with emergence time drawn form an exponential distribution, or to the mean of a population of cytonemes driven by a Poisson point process for their emergence. However, it is also of interest to characterize the variance or other statistics in the amount of morphogen cells receive. It turns out that when the cytonemes are independent of one another, regardless of how complicated the transport, attachment, detachment, and degradation processes are, the end result is that the distribution of morphogen received by cells is Poisson-distributed and thus the mean and variance are equal. We omit the details here, but the interested reader can find a description of this result in the Appendix.

### 3.2 Repeated packet delivery by attached cytonemes

Thus far we have only obtained exact results while assuming a single bolus of morphogen is carried by each cytoneme. But as we indicated earlier, in some cases sustained morphogen transport occurs along the length of a cytoneme in discrete packets called puncta for PIT or on receptors expressed on the surface of the cytoneme in RIT (*cf*. Figure 1). The primary effect of this alternative transport mechanism is that the amount of morphogen transported by any cytoneme will depend upon how long the cytoneme remains attached, not merely whether or not an attachment is made. Since morphogen is not delivered in one event, but distributed over time in a series of potentially-smaller events, longer connection time should be correlated with larger morphogen delivery. This contrasts with the bolus-based mechanisms, since the longer attachment time in that case merely delays the cytoneme return in the RIT mode, and has no effect in the PIT mode. For the punctum-based delivery mechanisms, the velocity of cytoneme extension as compared with punctum movement along a cytoneme also matters, and there are two cases, *v*_*p*_ ≤ *v*_*c*_ and *v*_*p*_ *> v*_*c*_. We begin with *v*_*p*_ ≤ *v*_*c*_ since it is easier, and then discuss the modifications needed for the latter case.

Consider a single cytoneme, formed as in the previous PIT model, and a single punctum which will be loaded onto the cytoneme sometime after the cytoneme is generated. For the RIT model with a punctum, transport along a cytoneme can be modeled as in the approach explained below, plus an additional time delay to account for the cytoneme extension phase from a receiver to a producer cell. In light of this similarity, we do not consider the RIT mode of punctum-based transport further.

We assume that the extension/retraction and attachment/detachment of a cytoneme in PIT mode are not affected by the punctum motion and have been described previously, and therefore we focus on a description of the punctum. To describe their motion, assume that the cytoneme is generated at *t* = 0, and let 𝒯_*h*_ be the time at which the cytoneme reaches a target cell and halts. The punctum is loaded into the cytoneme at a random time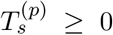, which is the delay, if any, from when the cytoneme is formed to when the punctum is loaded. We assume that loading is a Poisson process with parameter *λ*_*p*_, and note that there is no requirement that 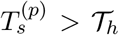 – the punctum can be loaded before or after the cytoneme attaches to a receiver cell. To simplify the analysis and results, we first condition the model on a fixed cytoneme length *x*_0_, and later generalize the result to arbitrary *x*_0_. Given *x*_0_, the punctum must arrive at its destination cell at a random time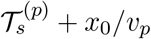, which accounts for punctum generation plus travel time along the cytoneme.

For a given *x*_0_, the governing equations for the puncta motion, puncta loading at *x* = 0, and morphogen reception at the receiver cell, are

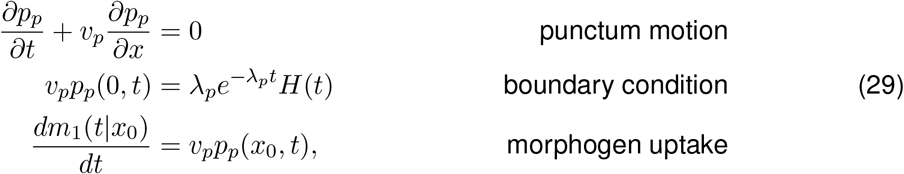

where *p*_*p*_(*x, t*|*x*_0_, 0) is the probability of the punctum being located at (*x, t*) ∈ (*x*_*o*_, *t*) given that it loaded at 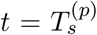 and that the cytoneme is attached to a cell at *x*_0_ . Since we assume that the punctum loading rate and transport velocity are independent of the cytoneme length, the full distribution *i*.*e*., not conditioned on an *a priori* value of *x*_0_, is found by multiplying *m*(*t*|*x*_0_) *× P*_*a*_(*t*|*x*_0_), the probability of a cytoneme being attached at position *x*_0_ at time *t*.

With *v*_*p*_ ≤ *v*_*c*_, the resulting solution for *m*_1_ is

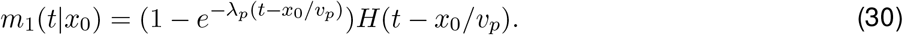

To understand how this leads to a distribution of morphogen throughout a tissue with multiple cytonemes and potentially multiple puncta on each cytoneme, we must now account for multiple puncta per cytoneme, and random cytoneme initiation times.

First, when multiple puncta are present, they are loaded onto the cytoneme at random times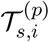. If we assume that these times are generated by a Poisson process of rate *λ*_*p*_, then, the expected morphogen uptake by the receiver cell attached at the terminal end of the cytoneme is

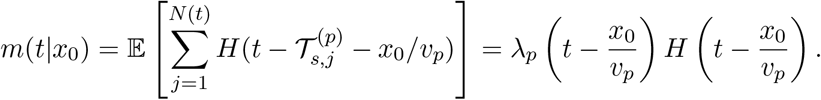

This result is linearly increasing in time after a time-delay needed for the punctum to reach the terminus of the cytoneme. Second, when the cytoneme start time is a random variable, the mean morphogen accumulation for a single punctum can be found by computing the expectation

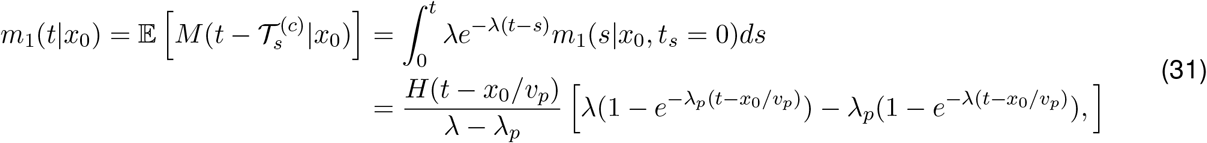

where 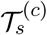 is the cytoneme start time, which we assume is exponentially-distributed with rate *λ*.

These results were conditioned on the cytoneme pausing at *x*_0_. In general, the pausing location is a random variable, *X* = *v*_*c*_𝒯^(*h*)^, which as in Section 3.1, is exponentially distributed as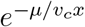. Since the pausing is independent of all of the other quantities in the model, the resulting distribution for *m*_1_ is found by multiplying the previous solution by this distribution, *i. e*.,

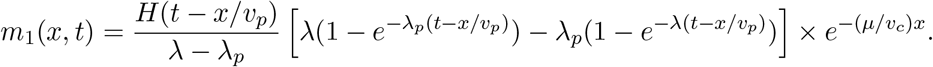

Combining multiple puncta with multiple cytonemes that are initiated at random times leads to a compound stochastic process for the cytonemes and puncta. Nonetheless, because the stochasticity is governed by Poisson processes, the model is still analytically solvable. In particular, the mean morphogen accumulation is

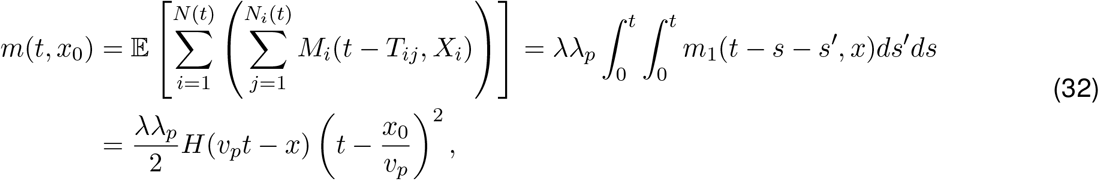

where *N* (*t*) is the total number of cytonemes, *N*_*i*_(*t*) is the number of puncta on the *i*th cytoneme, and *T*_*ij*_ is the generation time of the *j*th puncta on the *i*th cytoneme. The quantity *M* (*t*−*T*_*ij*_|*X*_*i*_) is a stochastic quantity representing the amount of morphogen delivered to a cell at 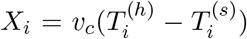 where 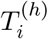 and 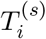 are the random starting and stopping time for the extending cytoneme, c.f. Section 3.1.

This result is quadratic since the morphogen accumulation is proportional to the number of cytonemes and the number of puncta. However, it is also unrealistic because this model supposes that cytonemes last forever. In reality, cytonemes extend, attach, and eventually retract. Thus, they only remain attached to receiver cells for a finite amount of time, limiting how much morphogen can be transferred by any individual cytoneme.

### 3.3 The effect of finite attachment times

Since the attachment times in the PIT-mode are finite, we stipulate that they are exponentially-distributed, and in this case the equation for *P*_*a*_(*x, t*) becomes

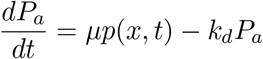

where *k*_*d*_ is the detachment rate and *µ* the attachment rate for the cytoneme to a receiver cell in the PIT mode.

Furthermore, morphogen can only be delivered if it has sufficient time to reach the delivery point after being loaded onto a cytoneme at the producer cell. This involves several additional modeling steps, and to simplify the analysis, we again begin by conditioning on a specific stopping length, *x* = *x*_0_, and assume that the cytoneme starts at *t* = 0 rather than at a random time.

This conditioning modifies the equations for *p*_*c*_ and *P*_*a*_ to become

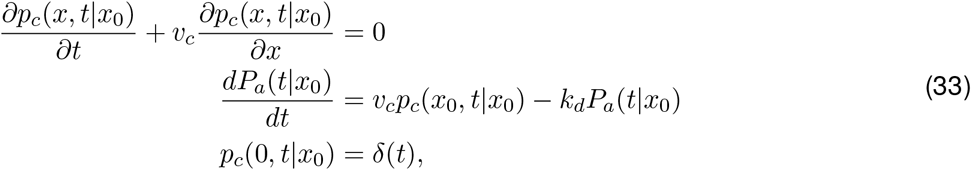

from which one obtains has solution

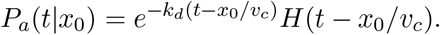

Since the time between attachment and detachment is exponentially-distributed, the probability density of detachment per unit time is then *k*_*d*_*P*_*a*_(*t*|*x*_0_). Hence, the probability, *P*_*d*_(*t*|*x*_0_) of detachment by time *t*, is

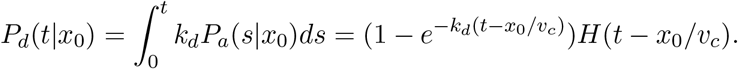

Given that the attachment time at *x*_0_ is exponentially-distributed, there is a non-zero probability that the cytoneme detaches before the punctum is delivered. This appears as a reaction rate −*r*(*t*|*x*_0_)*p*_*p*_(*t, x*_0_) per unit time in the partial differential equation for *p*_*p*_, and can be understood as the product of the probability of the cytoneme detaching at time *t* while the puncta is at position *x*. For the reaction rate *r*, we require the probability density for detachment in interval (*t, t* + *dt*), given survival up to *t*, or^8^

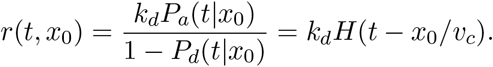

The model for a punctum now becomes

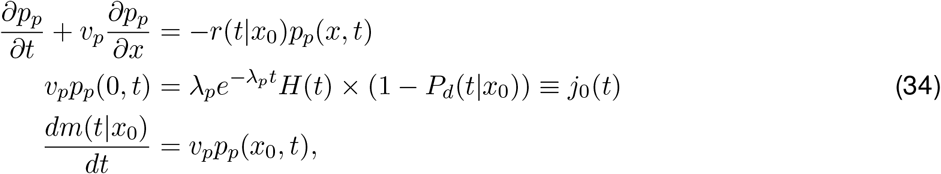

and the solution is for the expected amount of morphogen delivered is

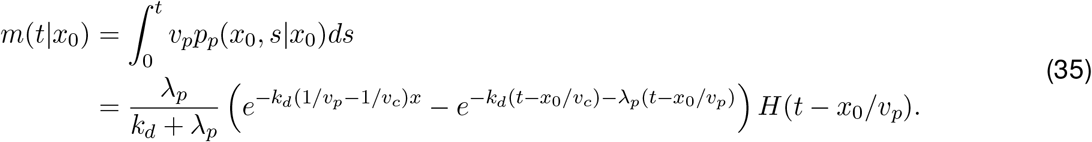

#### Remark

*For the more general advection-reaction equation p*_*t*_ + *vp*_*x*_ = −*r*(*t*)*p with boundary condition p*(0, *t*) = *P*_0_(*t*), *the general solution is*

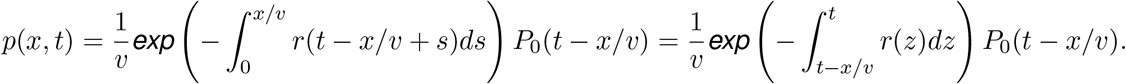

To demonstrate the validity of these results, we do a simple comparison of the analytical solution with a stochastic simulation. With the cytoneme conditioned to be generated at *t* = 0 and to attach to a cell located at *x*_0_, the only stochastic components are the detachment time, 𝒯_*d*_, and the puncta generation time, 𝒯_*p*_. The time for the puncta to arrive at the receiver cell is 𝒯_*p*_ + *x*_0_*/v*_*p*_. Thus, to simulate this process, we draw 𝒯_*d*_ and 𝒯_*p*_ from their corresponding distributions and observe whether 𝒯_*p*_ + *x*_0_*/v*_*p*_ *<* 𝒯_*d*_. After many trials, the results can be compared with the exact result for *m*(*t*|*x*_0_), as shown in Figure 7.

**Figure 7:**
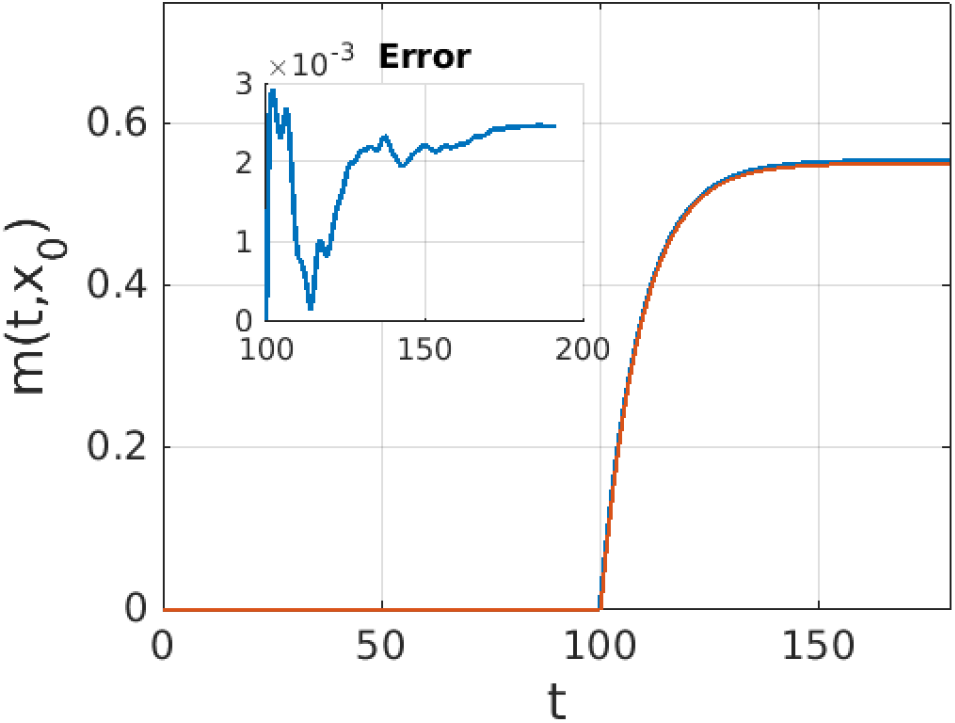
The comparison of simulation results with exact results for the repeated delivery mechanism. In this case, one punctum is generated randomly, and we observe the morphogen accumulation over 10,000 trials. The error is well within 1% upon comparison. Accumulation commences at *t* = 100 because of the time delay that cytonemes take to reach their target cell.

To obtain the unconditional dependence on *x*_0_, the solution for *m* can simply be multiplied by 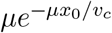 which is the probability density of halting at *x*_0_ = *x*. Likewise, to remove the *t* = 0 condition, the solution can be convolved with *ψ*(*t*) = *λe*^−*λt*^, the distribution for the cytoneme starting time. In the limit *k*_*d*_ → 0, the earlier results are obtained, and in addition, the morphogen accumulation for arbitrary *k*_*d*_ at large times is

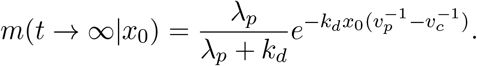

Other results left for the reader to verify are (i) the probability to not generate a punctum prior to detachment (*λ*_*p*_*/*(*λ*_*p*_ + *k*_*d*_)), and (ii) the probability for detachment of the cytoneme after the punctum has been loaded, but before it arrives at the distal end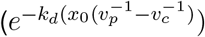.

When *v*_*p*_ *> v*_*c*_, the approach is very similar except that we have to separately track the probability for the punctum to reach the leading edge of the extending cytoneme prior to attachment. Conditioning as before on *x*_0_ and *t* = 0 to start, the model is essentially the same as before, except that now the time-delay factor for morphogen accumulation to commence is *x*_0_*/v*_*c*_ rather than *x*_0_*/v*_*p*_. To compute the probability of the punctum reaching the tip of the cytoneme when *v*_*p*_ *> v*_*c*_, define the time, Δ*t* = (*v*_*p*_ − *v*_*c*_)*/x*_0_ where a punctum created before Δ*t* will reach the cytoneme tip prior to attachment. The cytoneme tip position is given by *x* = *v*_*c*_*t*, and hence the accumulation at the tip is

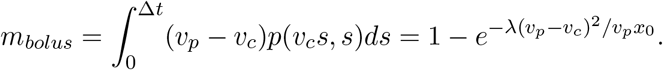

The full solution is now this probability plus the previous solution for the continual transport of the punctum along the cytoneme.

Finally, these results are easily extended to multiple puncta traveling along a cytoneme. In this case, the stochastic process is a compound Poisson process. The puncta are assumed to be generated at random times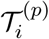, and this process is conditional on the cytoneme existing, which in turn is governed by a stochastic process.

To see how this works, consider the PIT model with a single cytoneme generated at *t* = 0, but multiple puncta. The cytoneme extension is the same as earlier, but now, the equation for the puncta is governed by a boundary condition of the form

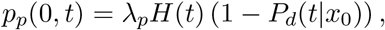

where *λ*_*p*_ is the rate of puncta formation. The solution in this case is of the form

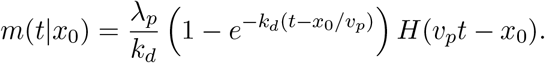

This leads to an average of *λ*_*p*_*/k*_*d*_ units of morphogen delivered per cytoneme.

If now, we suppose that there are multiple cytonemes being generated, starting from *t* = 0, at rate *λ*, we can compute the expected morphogen amount via

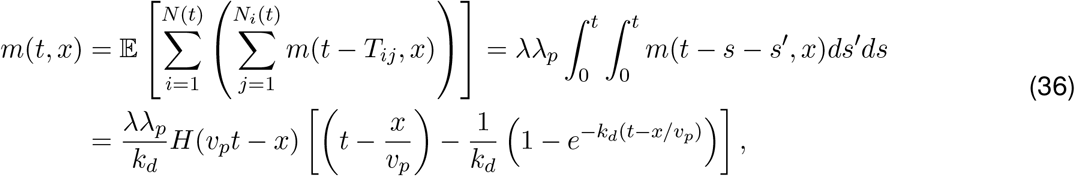

where *N* (*t*) is the total number of cytonemes, *N*_*i*_(*t*) is the number of puncta on the *i*th cytoneme, and *T*_*ij*_ is the generation time of the *j*th puncta on the *i*th cytoneme. In the limit *k*_*d*_ → 0, this reduces to a finite limit,

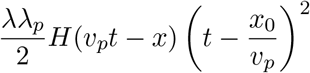

which is quadratic since the morphogen transport is proportional to the number of cytonemes and the number of puncta per cytoneme.

Modeling each punctum is a high level of detail, and in some limits, such detailed models are not necessary. For instance, assume that puncta are formed much more rapidly than cytonemes, and that a typical cytoneme remains attached to a target cell for a long time compared to the time to transfer a single puncta. The result is that when a cytoneme attaches to a target cell, a large number of puncta are transferred at essentially a continuous rate. This allows us to assume that the amount of morphogen transferred is proportional to the length of time the cytoneme remains attached. In this case, model for the puncta transport in the PIT model reduces to the following:

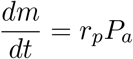

where *r*_*p*_ is the morphogen transfer rate from the puncta, and *P*_*a*_ is as before, the probability of being attached at time *t*. In this case, the amount of morphogen transferred is

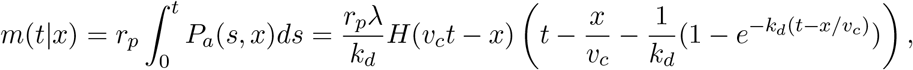

which reduces to equation (36) with *λ*_*p*_ replaced by *r*_*p*_ and *v*_*p*_ replaced by *v*_*c*_.

For the RIT-mode with morphogen puncta, the same approximations can be applied. The calculations are more tedious, but results are attainable,

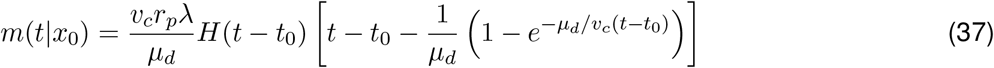

with

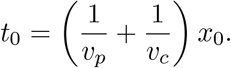

In each case, the result is similar - an initial period of no accumulation followed by gradual ramp up to a linear rate whose slope depends upon the cytoneme velocity, production rate, and the amount of morphogen per puncta.

#### Remark

*While a given cytoneme may well undergo multiple periods of attachment and motion to reach multiple cells, we have not modelled this possibility. However, it is important since the resulting distribution of morphogen could be identical if one cytoneme makes multiple attachments at different cells, or if multiple cytonemes from the same cell make single attachments. It is possible that it would take a cell less effort to produce a single cytoneme that makes multiple attachments than to generate many cytonemes, and we believe that this idea would be worth exploring experimentally to determine whether such behavior occurs, and what processes set the balance between multiple cytonemes making fewer stops – leading to more exploration of the local environment– or fewer cytonemes making many stops that may lead to more morphogen transport*.

## 4 Stochastic simulation of RIT transport

While useful insight can be gained from the simple models above, if more complicated or non-linear interactions between cytonemes and cells are important, then simulations may be necessary. Here we focus on the more complicated RIT-transport mode, but PIT can be studied in the same framework described below.

The simulation approach is straight-forward since it is relatively easy to develop effective numerical methods for cytoneme dynamics. The high-level approach is essentially an agent-based model with Monte-Carlo type steps that govern changes in the states (e.g. from resting to motile) of each cytoneme. Furthermore, even complex phenomena such as cytoneme-cytoneme interactions, which would be difficult to describe via the simple evolution equations in the previous models, can be simulated in a straightforward manner with the numerical methods. The numerical methods are based on discretization of the evolution equations for the cytoneme location, e.g.

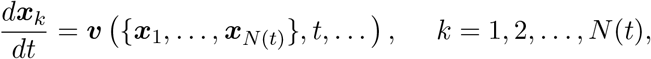

and also incorporate additional modules to describe cytoneme attachment, generation, and morphogen transfer along cytonemes. In contrast to equation (7) for Newton’s Law of motion, we assume the velocity is constant as we have elsewhere, but in principle a very similar stochastic simulation approach could be used when the velocity is also a stochastic quantity. To our knowledge, the use of a stochastic cytoneme-level simulation of morphogen transport is novel. The closest previous effort we are aware of isthe modeling in effort in (Aguirre-Tamaral & Guerrero 2021) which does incorporate some aspects such as cytoneme motion, such as the cytoneme length in the model. However, their model is ultimately still formulated at a continuum rather than stochastic level.

### 4.1 Comparison of analytical results for RIT with results from computational stochastic models

To begin, we use a simple stochastic simulation procedure described above and validate the results of simulations by comparing them with the analytical results in the previous sections. In Figure 8, results from the RIT model based on the processes shown in Figure 10(a), as well as results obtained from averaging 100 stochastic simulations (dots), both in one space dimension, are plotted at several times. The maximum error tends to occur near the leading edge of the morphogen distribution, and tends to be no more than approximately 0.2 morphogen-packets/cell.

**Figure 8:**
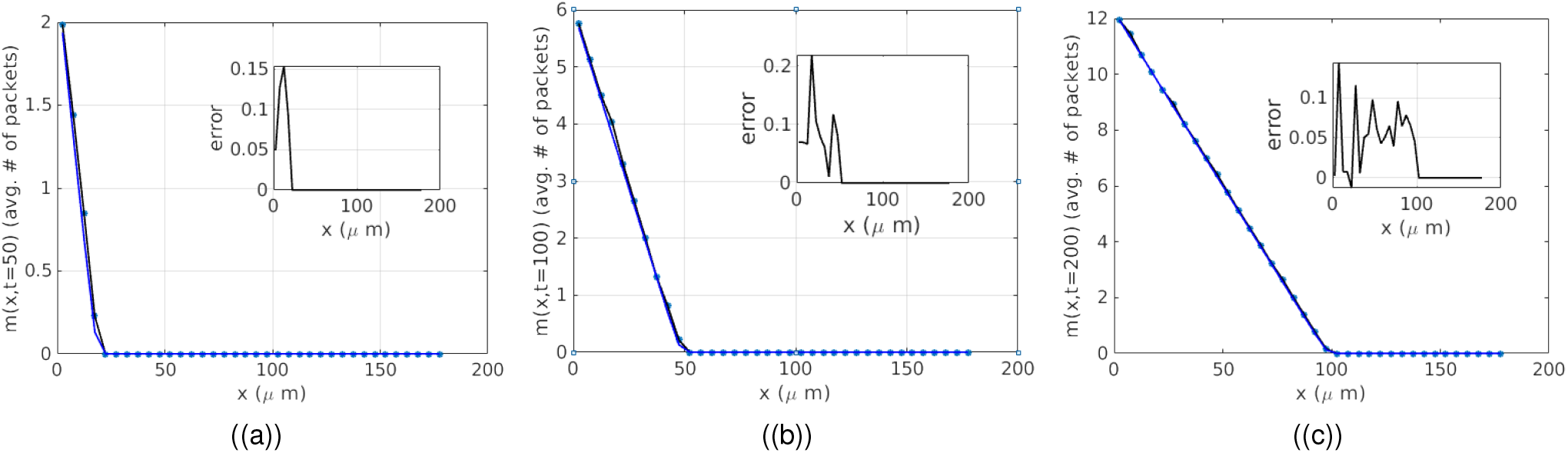
Comparisons of *m*(*x, t*) between simulations and analytical results at *t* = 40, 100, and 200 minutes (left to right). Shown are the number of packets per receiver cell as a function of distance from the source at x = 0.The maximum error is roughly 0.2, and tends to occur at the edges of the morphogen distribution, probably due to small cytoneme numbers at the edges, and possibly from small numerical errors in determining when the cytonemes start or stop. In these simulations, *µ*_*d*_*/v* = 1 and *λ* = 0.25/min.

Under assumptions discussed earlier and using results from the appendix, the distribution of the amount of morphogen received should be Poisson-distributed in time, with a rate determined by the first passage-time distribution for morphogen delivery. We can also check that this is the case by analyzing the distribution of received morphogen for cells at various distances from the producer cells. Several such choices are shown in Figure 9. A characteristic of Poisson distributions is that their mean and variance are equal, and the computational results reflect this, as shown in 9(a). Similarly, the distribution of *M* (*t*|*x*_0_) at two values of *x*_0_ are shown in (b) and (c). The blue bars represent simulated data, and the black curves are analytical results for the distribution of morphogen at *x*_0_. In (b) and (c), there is still some error between the observed distribution and the expected distribution. This noise disappears at larger time as more morphogen is accumulated, but it is difficult to evaluate the factorial and exponential in the Poisson distributions for large values of the mean, thus a smaller time (*t* = 200*s*) is used for (b) and (c) than (a) which is for *t* = 1000*s*.

**Figure 9:**
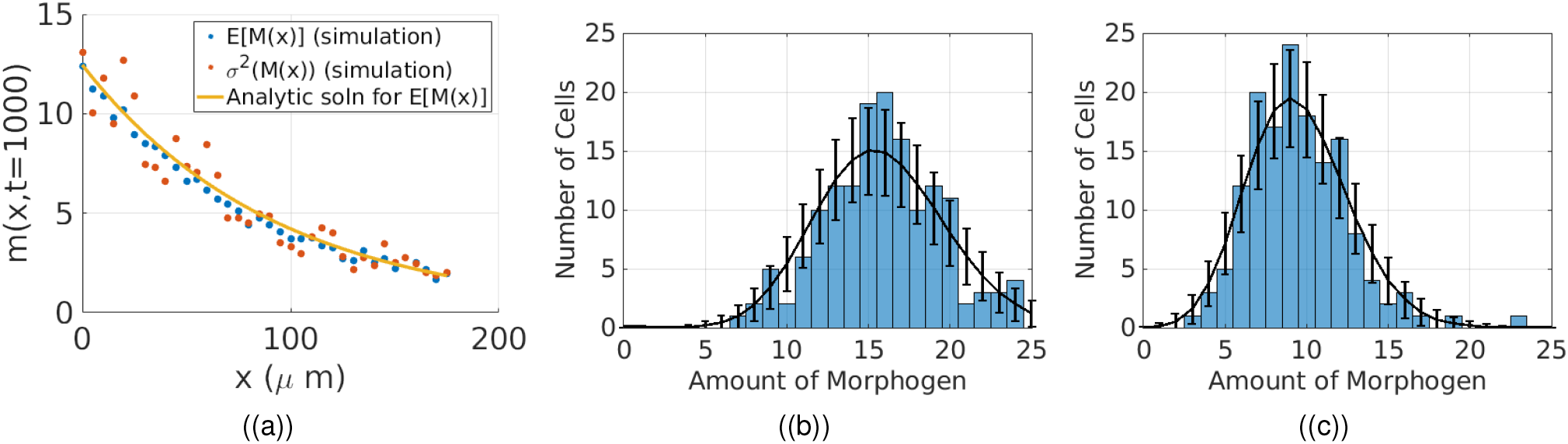
**(a)** The analytical (yellow curve) and computational mean and variance (blue and red dots). In **(b)** and **(c)** the distribution of morphogen uptake for receiver cells centered along a line of fixed *x* at *x*_0_ = 42.5*µm* and *x*_0_ = 62.5*µm* resp., are shown and compared with the expected Poisson distribution (errors bars are *±* one standard deviation from the mean).

### 4.2 Extensions to the cytoneme models

Now that we have compared the simplest stochastic models and found agreement with the analytical results we can take advantage of the fact that it is relatively easy to include various biological effects in the stochastic simulation algorithm. The tissue geometry used consists of an array of hexagonal cells as shown in Figures 10(b,c), which show typical computational results. The parameters for a typical solution are listed in Table 1.

**Figure 10:**
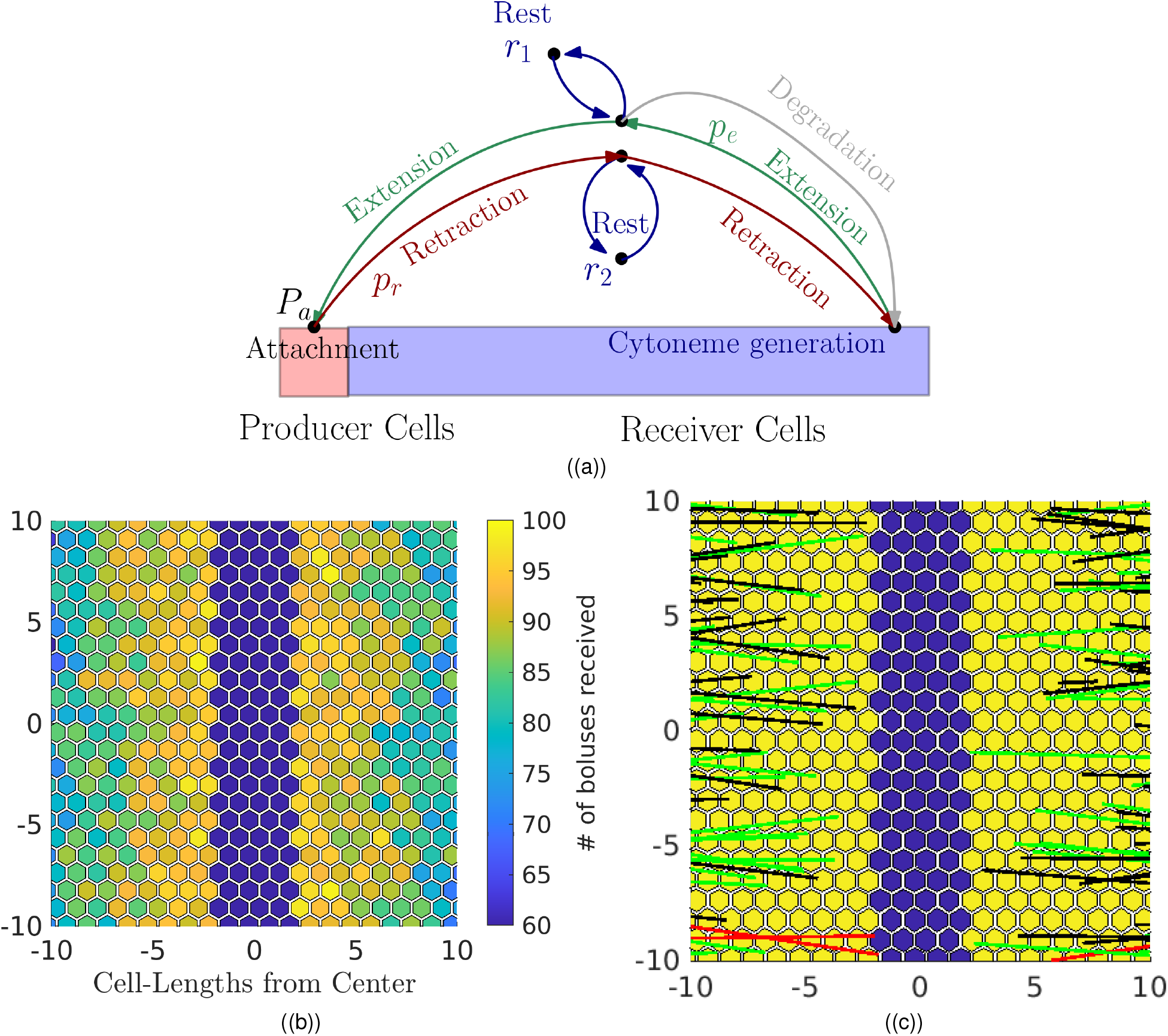
(a) Diagram of the various transitions that can occur in the RIT model. Cytonemes start in the extension phase after generation. At any point, they may enter a rest state at a rate *λ*_*r*,1_, or spontaneously begin retracting without reaching the source (degradation). If they reach the source, they become attached, and then undergo retraction, or enter intermittent rest phases at a rate *λ*_*r*,2_ as they retract. (b) Depiction of the result of a typical simulation. The coloring corresponds to the number of morphogen packets received by each cell and the colorbar gives the number scale. The four central columns of ’blue’ cells are the producer cells. Their morphogen content is not counted since they produce rather than receive morphogen. The length scale on the axis is cell diameter (∼ 10*µm*), hence *x* = 2 corresponds to a distance 20*µm*, or 2 cells from the center. (c) Diagram of the cytonemes during a simulation. Black cytonemes are extending, red are retracting without morphogen, and green received a morphogen packet and are retracting.

**Table 1:** Typical parameter values for the stochastic simulation method.

| Symbol | Definition | Default Value |
| --- | --- | --- |
| $\lambda$ | Cytoneme generation rate | $0.015\text{s}^{-1}$ |
| $\mu$ | Cytoneme attachment rate | $3\text{s}^{-1}$ |
| $v_0$ | cytoneme velocity | $0.1\mu\text{m/s}$ |
| $k_d$ | cytoneme failure rate | $0.01\text{s}^{-1}$ |
| $\theta$ | cytoneme angle | $-\pi/8 \leq \theta \leq \pi/8 \text{ rad}$ |
| $\delta t$ | time-step length | $1\text{s}$ |

First, cytonemes do not always travel directly towards their targets, and therefore we generate their direction according to an angular distribution. If *θ* = 0 is the optimal direction, we choose an initial angle from a uniform distributions *θ* ∈ [−Θ, Θ] and maintain this direction throughout movement.

Theoretically these results can also be obtained analytically if cytonemes travel in a straight line at an angle Θ. In this case, the FPT transfer of morphogen to a cell only depends on *θ* via the effect that *θ* has on the length the cytoneme must travel to reach a producer cell. Thus, the average rate of morphogen uptake is given by

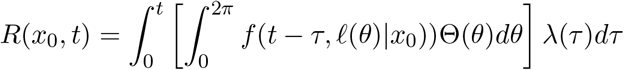

where we have assumed that the cytoneme angle and generation rate are independent, and governed by distributions Θ(*θ*) and *λ*(*t*) respectively. The inner term in the integral can be thought of as an angle-averaged FPT being convolved with the cytoneme generation rate. If the producer region is simply a stripe of cells in a line, we do not expect that the angle significantly alters the morphogen uptake results, except that the FPT must be averaged over the angular coordinate, which leads to some to some loss of efficiency when compared with cytonemes that travel directly to the closest prodcer cell. Nonetheless, as more becomes known about cytonemes, it may be important to take into account changes in direction as the cytoneme travels.

### 4.3 Modeling of morphogen puncta

We have considered cytonemes as the main component of the simulation methods, although cells are ultimately responding to uptake of morphogens. When cytonemes deliver morphogens as a single bolus, tracking the bolus size, and the rate at which cytonemes return to their host cell in RIT mode is sufficient for describing the morphogen distribution. However, morphogens are often transported along cytonemes as a series of discrete puncta, and not as a single bolus as discussed in Section 3.2. Here we extend the stochastic simulations to explicitly include the puncta of morphogens.

To this end we use a more complex stochastic simulation, one component of which is defined by the fact that the *internal* state of a cytoneme changes when the cytoneme changes state from extending to attached to a producer cell, and begins uploading morphogen. Upon attachment, we simulate the uploading using a Poisson process at constant rate, *λ*_*p*_ to determine times at which puncta are loaded onto the attached cytoneme. After one is loaded onto a cytoneme, we update its position by computing its velocity and using a forward Euler approximation in each time-step. When the puncta reaches the end of the cytoneme, we update the amount of morphogen in that cell according to the amount in each puncta. The results for a simulation are shown in Figure 11. Here we assume that each puncta simply contains one unit of morphogen-content, but this could also be a random variable as indicated in equation (7).

**Figure 11:**
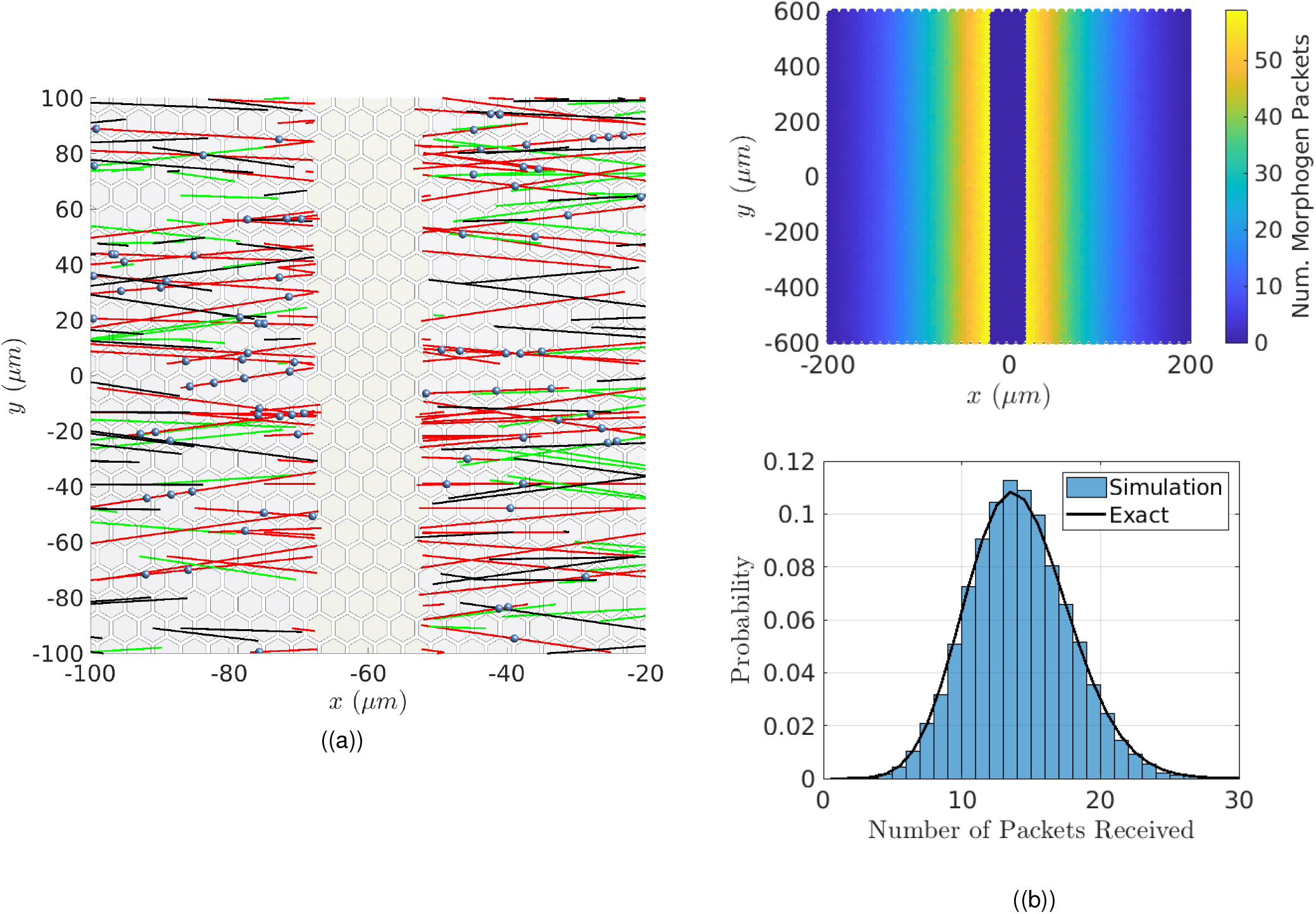
(a) Trajectories of the morphogen packets (blue spheres) moving along cytonemes in a portion of the domain. Green cytonemes are retracting, red are attached, and black are extending. (b) Morphogen accumulation, *m*(*x, t*) (top) and statistical variation (bottom) in cells located at *x*_0_ = 5.75 In (b-lower) the x-axis is the morphogen amount, and the y-axis the probability, or proportion of cells at *x*_0_ that receive a given quantity of morphogen. The results in (b) are averaged over 150 simulations.

## 5 Other generalizations

### 5.1 Morphogen-dependent feedback in the receiver cells

A significant advantage of the stochastic simulation method is that it is easier to include nonlinear interactions between components and effects that involve complicated probability distributions. Such interactions can be included in evolution equations for *p*(*x, t*), but the resulting equations are typically high-dimensional and it is difficult to obtain simple equations for the mean or standard deviation.

As another example of what can be investigated via stochastic simulations we consider a simple form of feedback, whereby a cell can detect its total morphogen load and down-regulate cytoneme generation when it achieves a sufficient amount. This model is plausible since cells are sensitive to the amount of morphogen received, and down-regulating the extension of cytonemes is one method of control. This leads to models of the form

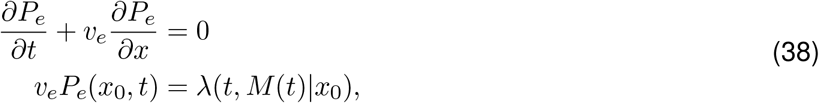

where *P*_*e*_(*x, t*) and *M* (*t*) are coupled via the cytoneme generation rate. However, it is difficult to extract statistics other than the mean from this type of model, since the resulting stochastic process is no longer a Poisson point process, or even a Markov process in this case. Nonetheless, it is straightforward to simulate this type of model once a choice for *λ*(*t, M*|*x*_0_) is specified.

As an example, let us assume that a cell gradually ceases cytoneme production upon receiving a sufficient level of a morphogen. In this case, we set *λ*(*t, M*) to be a decreasing function of *M*, and in the simplest case, *λ*(*t, M*) = *λ*_0_ − *λ*_1_*M* is a linearly-decreasing function. The ratio *λ*_0_*/λ*_1_ determines the maximum amount of morphogen the cell will receive before it shuts off cytoneme generation, and in the computations in Figure 12 this is set at 100. All cells eventually reach a value near this steady-state, and if we choose a value of *x*_0_ and plot the distribution of morphogen amounts, the feedback leads to a plateau-like distribution concentrated at the shut-off level. The establishment of the distribution also differs from the previous cases, in that in this case there is a wave-like propagation of the saturation plateau, as shown in Figure 12.

**Figure 12:**
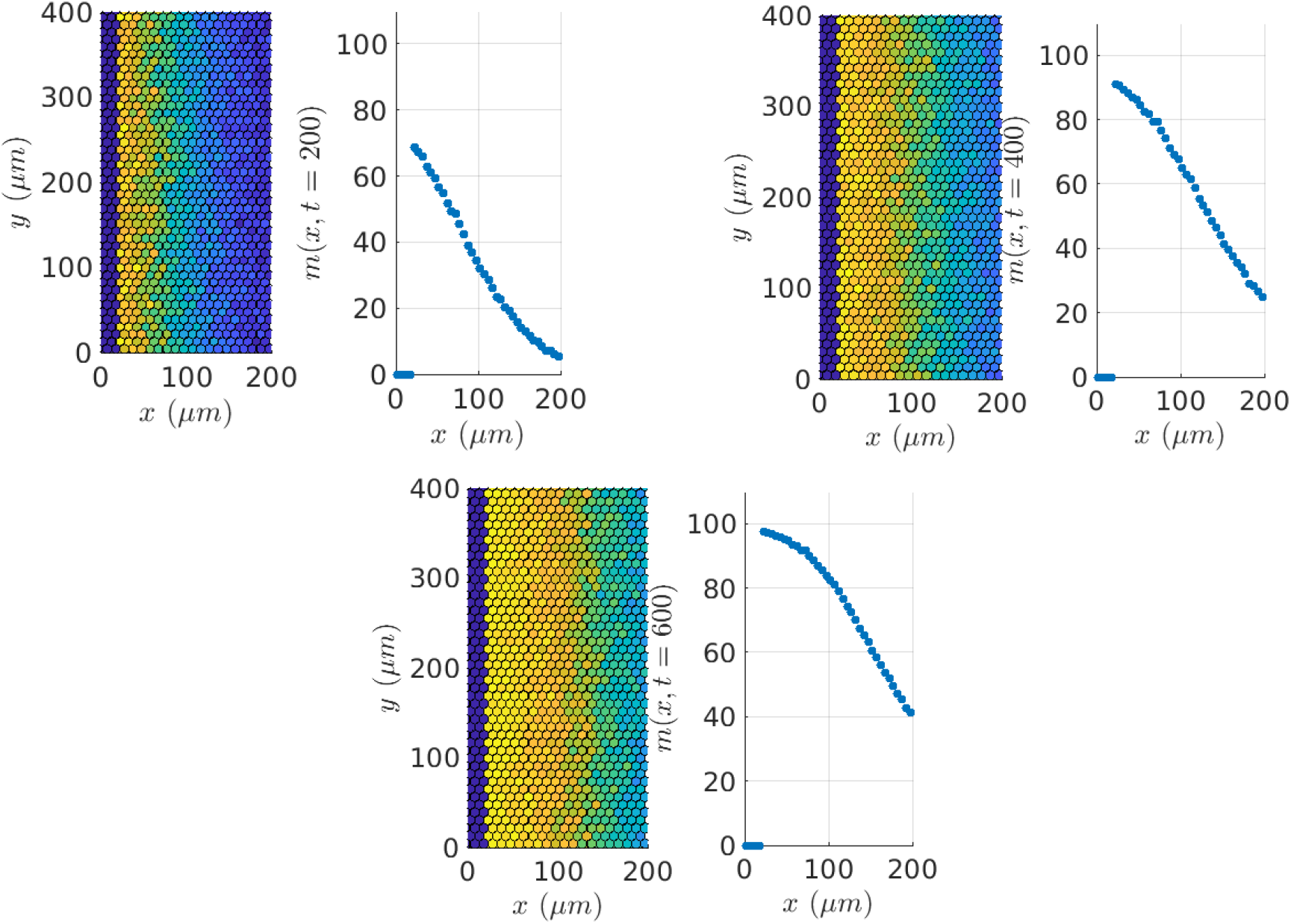
Evolution of the steady-state concentration as time increases. Left to right: *t* = 200, 400, 600*s*. The hexagonal arrays in each show the morphogen concentrations in each cell. The scatter-plots show the mean morphogen level as a function of *x*_0_. On the left the maximum level has not been reached by any cells, but in (b) and (c), the saturation level is reached by increasingly distant cells.

### 5.2 The effect of tissue growth on morphogen distributions

Another important generalization is when morphogen fields are established on similar time-scales with tissue growth. Throughout we have implicitly assumed that cells are growing slowly or not at all compared to the speed of cytoneme-transport. However, when growth and transport occur on similar time-scales, changes in cell position alter the resulting distributions of morphogen. In general tissue growth is dependent on the morphogen levels, and the problem becomes coupled and nonlinear. However, if we assume that it is independent of the morphogen, but patterning is still morphogen-dependent, then it is possible to derive analytical solutions for the morphogen accumulation.

Begin by defining a coordinate system in which the morphogen-producing cells remain fixed at *x* = 0, but receiving cells in the growing tissue can move relative to the producing cells. For instance, if the growth rate *γ* is spatially uniform and the tissue is one-dimensional, then the tissue velocity at any point is *v*_*t*_(*x, t*) = *γx* in a fixed coordinate system with *x* ∈ [0, ∞). Therefore the position at time *t* of a cell that was at *x*_0_ at *t* = 0 is the solution of

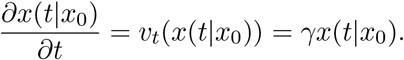

If *γ* = *γ*(*t*) is a function of time, then

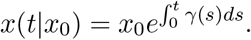

In early phases of growth *γ* is approximately constant, and the size increase is exponential. Eventually growth slows down and *γ* → 0 for large *t* as the final tissue size is obtained. On the other hand, if *γ*(*t*) = *β/*(1 + *βt*) for *β >* 0, then *x*(*t*|*x*_0_) = *x*_0_(1 + *βt*). We will use this choice as it is convenient as a starting case where exact solutions are easy to compute. It also can be thought of as a first-order approximation for small *t* of the exponential growth example since *e*^*γt*^ = 1 + *γt* + 𝒪 (*γt*)^2^.

In PIT the details of the cell structure are not needed. Since morphogen delivery is per unit length, we can simply let the length grow without quantizing it into cells. We do not have to change the parameters in the model that govern cytoneme dynamics, in particular the stopping time, if we assume that the cytoneme extension does not depend on the underlying tissue. All that is needed in this case is the distribution of cytonemes, in particular the distribution in any interval (*x, x* + *δx*) and from this one could extract the amount of morphgen received in any time interval and in any space interval. This avoids having to define what a ’cell’ does, and trying to include divisions. Since the cytoneme halting rate is assumed to be unchanged by the growing tissue, the rate at which cytonemes halt per unit length is then *µp*(*x*(*t*), *t*)*J*(*x*(*t*), *t*), where *J*(*x*(*t*), *t*) is the Jacobian of the map *x*(0) *1*→ *x*(*t*). Thus the morphogen accumulation is now governed by

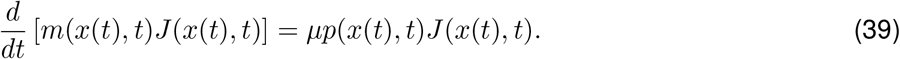

This formulation is needed because the growing cells are not passive particles that move along a trajectory in space, in which case integration of *µp*(*x, t*) along each trajectory would be sufficient. Rather, the cells and tissue grow over time, and this must be accounted for by the Jacobian as above. This also ensures that *m*(*x, t*) maintains its definition as an amount of morphogen per unit length in the tissue as it grows. This result is a natural generalization of earlier results, because when there is no motion *J*(*x, t*) = 1.

If we use the growth law *x*(*t*|*x*_0_) = *x*_0_(1 + *βt*), the solution of equation (39) is

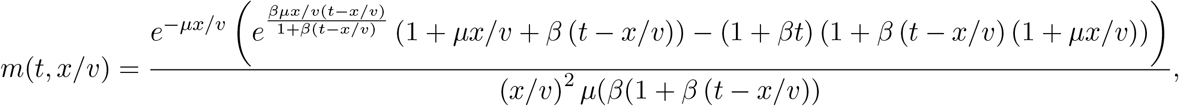

where *v* is the cytoneme extension velocity. This solution approaches the solution given earlier at equation (26) (with *λ* = 1) in the limit as *β* → 0, and the leading order terms are

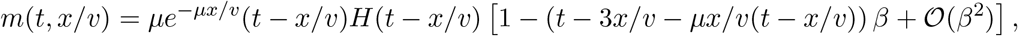

which is accurate for small *β*. In this result, there is a tradeoff between higher growth and morphogen transport. As the growth rate increases, the morphogen distribution becomes broader as seen in Figure 13, but less morphogen is delivered to nearby cells as *β* increases.

**Figure 13:**
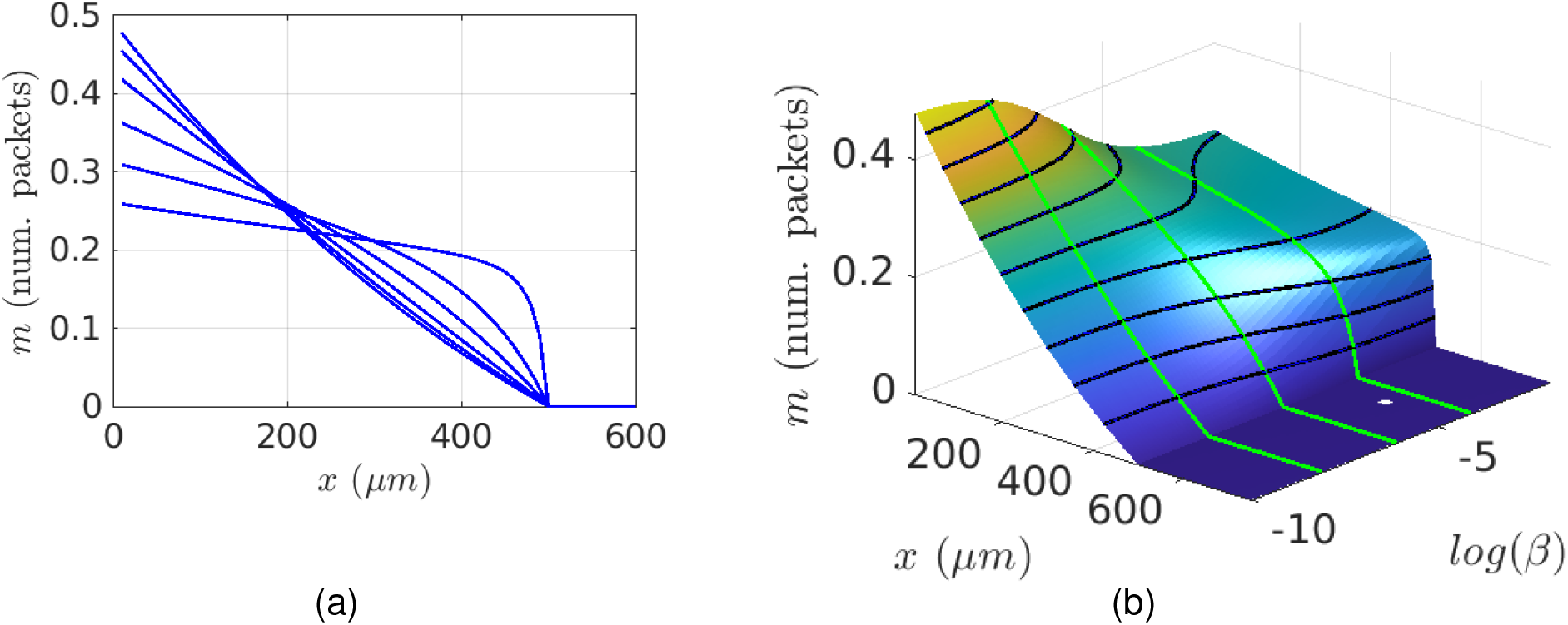
(a) The morphogen distribution at *t* = 500 for several values of the growth rate, *β*. (b) Surface for *m*(*x, t*|*β*) with *t* = 500 and *log*(*B*) in the range (−10, −2). Green curves indicate *m*(*x, t*) for several fixed values of *β* highlighting the change in the shape of the distribution. Black contour lines are solution sets of *{m*(*x*(*β*), *t* = 500|*β*) = *m*_0_} with *m*_0_ ∈ {0, 0.05, 0.1, …, 0.5}.

For large *β*, a finite limit also exists and is equal (up to 𝒪 (*β*^−1^)) to

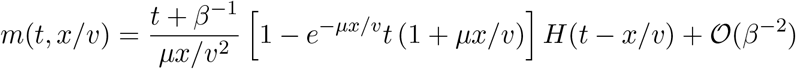

and we see that the leading edge of the distribution eventually becomes discontinuous since in the limit *x/v* → *t* from below, *m* = (*µt*)^−1^(1 − *e*^−*µt*^(1 + *µt*)).

When the growth rate *γ* is a constant, then *x*(*t*|*x*_0_) = *x*_0_*e*^*γt*^, and the solution takes the form

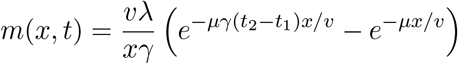

where *t*_1_ and *t*_2_ are the first and second real-valued solutions (when they exist) of the transcendental equation,

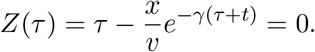

For *t < t*_1_, the result is 0, and when *t*_1_ *< t < t*_2_, set *t* = *t*_2_. While the transcendental equation has a solution in terms of the Lambert-W function, the details are more complicated due to the need to distinguish between the first and second solutions. We omit the details, but note that for small *γt*, the results appear to be qualitatively very similar to the preceeding results, which is anticipated by the fact that a Taylor expansion of *e*^*γt*^ in *γ* is 1 + *γt* up to order (*γt*)^2^.

## 6 Diffusive vs CBT in a complex tissue environment

Now that we have discussed the CBT modes at length, we turn to the original question that motivated this work: how do diffusion and CBT mechanisms compare? Both the Turing and positional-information mechanisms are based on diffusion in a homogeneous medium. The latter posits a mechanistic description of the form

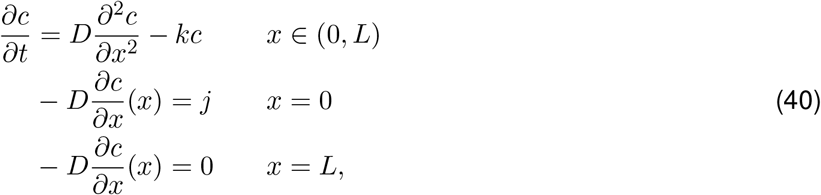

for establishment of the morphogen profile, but this rarely suffices to describe the detailed processes that are involved in biological tissues. For example, extracellular signals are detected by receptors, and this simple process adds terms for binding and release from the receptor to (40), as well as a separate equation for the binding and release at the cell (Lander *et al*. 2002).

An example of a complex tissue in which patterning occurs is the *Drosophila melanogaster* wing disc mentioned earlier and shown in Figure 14. The disc has two cell layers separated by a fluid lumen (Figure 14(c)), one a layer of columnar epithelial cells with the apical side at the lumen and a peripodial epithelium overlying the lumen (Gibson *et al*. 2002)(Figure 14(c)). The lateral membranes of adjacent cells are connected via two classes of junctions – ZAs and SJs – that separate the extracellular fluid into apical and baso-lateral layers (Gibson & Gibson 2009; Harris & Tepass 2010; Choi 2018) (details of the entire developmental process are given in (Gou *et al*. 2020)). Of relevance here is the fact that this is a tissue in which diffusion and cytonemes are thought to play roles in the transport of the morphogen Decapentaplegic (Dpp). Hence it is an ideal example in which a comparison between diffusion and cytoneme-based mechanisms could shed light on their respective roles in development.

**Figure 14:**
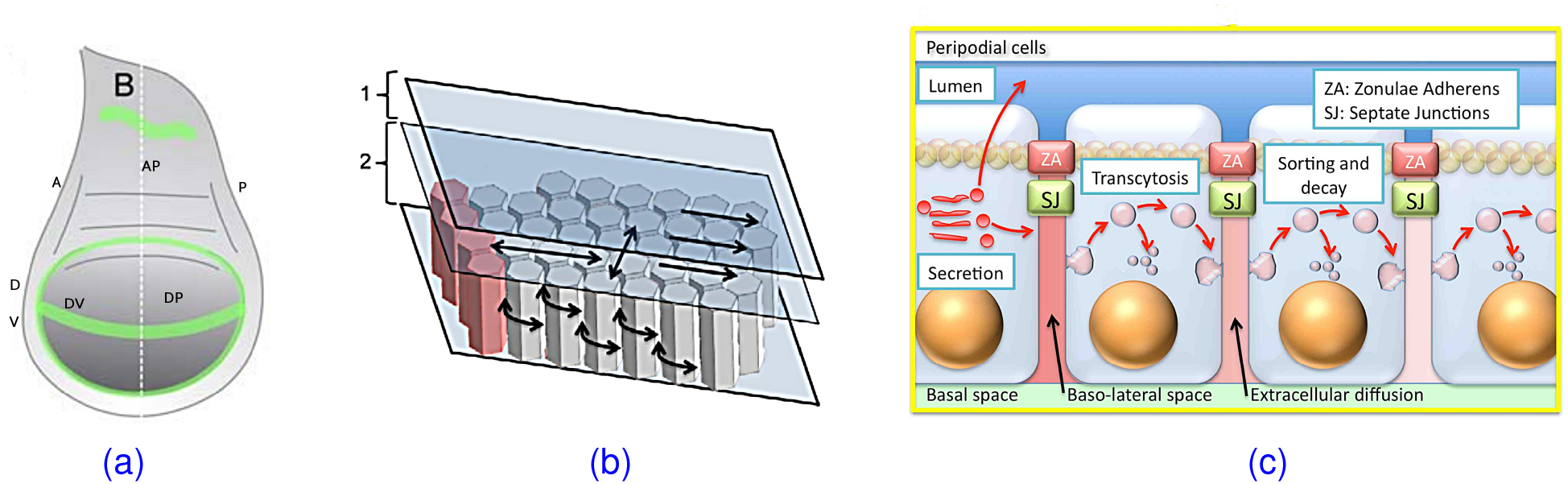
(a) The wing disc with the disc proper (DP), and anteroposterior (AP) and dorsoventral (DV) axes indicated. (b) A section of the disc, showing the fluid luminal section (1), the hexagonal cells below (2), and the source of Dpp at the left along the AP boundary in (a) From (Umulis & Othmer 2013). (c) A vertical cross-section showing the transport processes that affect the morphogen distribution (From (Gou *et al*. 2020).)

### 6.1 From micro-to macro-parameters for diffusion and reaction

In a previous paper (Stotsky *et al*. 2021), hereafter referred to as **I**, we analyzed how microscale processes that are involved in transport in complex tissues are reflected in the macroscopic diffusion and decay terms in a system such as (40). To frame the comparison with cytoneme transport, we briefly describe some of the underlying ideas in that work. The interested reader can find details of the underlying analysis in **I**.

The diffusion based models in **I** are based on continuous-time random walks (CTRWs) along edges of a network consisting of edges and/or centers of cells. In these models, molecules such as Dpp can be transported between adjacent cells via a number of steps (diffusion, binding, etc.). Starting at a subcellular level, it is typical to assume that the individual steps occur with exponentially-distributed waiting time distributions, and it is then possible to obtain the first-passage-time (FPT) distribution for a molecule to move from one cell to the next. Once the FPT is known, one can extract macroscale equations that govern the resulting system at large length and time scales. The macroscale equations are obtained by considering the limit as the size of an individual cell (*ℓ*) becomes very small compared to the size of the tissue (*L*), and the ratio *ϵ* = *ℓ/L* serves as an expansion parameter from which the leading order terms give the macroscale equation. The resulting macroscale equation is typically a diffusion-advection-reaction equation whose parameters in turn reflect the details of the underlying cell-level model. A simple example highlighting the details of the method is given in Appendix 8.6. Here we focus on a more complex example in a hexagonal lattice and use this as our basis for comparing cytoneme-based and diffusive transport.

#### 6.1.1 Diffusve transport

The wing-disc, as is typical of many epithelial tissues, consists of tightly-packed cells that can be approximated as having a hexagonal cross section as in Figure 14(b), and the 2D lattice shown below can be thought of as a horizontal slice through the tissue shown there.

As is shown in Figure 16(a), regular hexagonal lattices can be constructed as an alternating pattern of two topologically-distinct sets of points. The lattice is called a honeycomb or Bravais lattice, and the distinction between points stems from two facts. Firstly, all points are connected to three neighbors, and each point has two lateral connections, but the blue (circle) points connect upwards to a red (square) point, whereas the red points connect downwards towards a blue point. Thus, the red and blue points are topologically-distinct because they exhibit a mirror-image symmetry, but not translational symmetry under translations by an edge length. It is important to note that this partitioning of the points stems from the topology of the hexagonal lattice, and does not imply that different dynamics exist at the Type I and Type II points; the waiting-times and all other physical properties associated with the Type I and Type II points are assumed equal in the model.

Since the hexagonal lattice has two distinct types of points, we have to consider jumps on an edge from Type I to Type II and from Type II to Type I – there are no direct jumps along edges that take a particle from a Type I point to another Type I point. In effect the edges of the graph are ’directed’. Let *T*_12_ denote the transitions from Type II to Type I (see the shaded triangle labeled 𝒯 in Figure 16(a)), and let *T*_21_ denote transitions from Type I to Type II. Then the equations for the occupation probabilities take the form

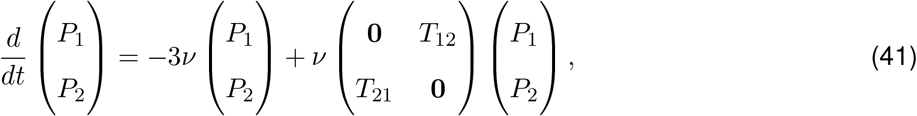

where *ν* is the transition rate. The transition operators, *T*_12_ and *T*_21_ are defined as

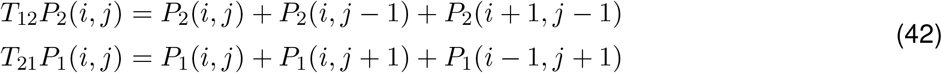

and are in fact adjoints of one another, *i. e*.,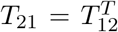. With the alternating structure delineated in Figure 16(a), it is convenient to solve for the Type II point probabilities in terms of the Type I points to obtain a single equation for the Type I points. For simplicity, assume that the initial condition is concentrated on Type I points only, the general case is not difficult to obtain but involves more book-keeping which obfuscates the underlying method. This leads to a new equation of the form

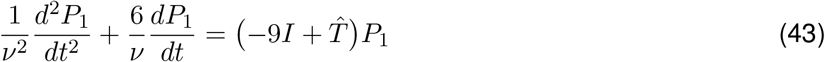

with the new transition matrix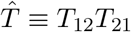.

The operator *T*_12_*T*_21_ can be found by computing

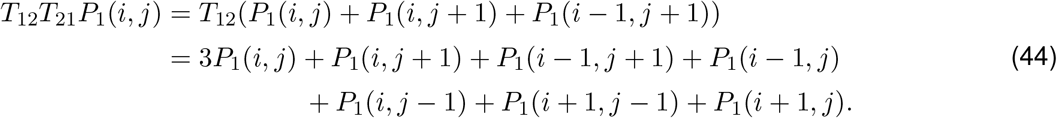

From this, we see that the right-hand side of equation (43) is just Δtri*P*_1_, the discrete Laplacian for vertex-to-vertex jumps on a triangular lattice^9^. Compared with the hexagonal lattice with edge-length *ℓ*, the triangular lattice in this case has edge-length 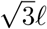 since the resultant length of any two adjacent edges of the hexagonal lattice is 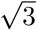 times the length of the individual edges (see Figure 15(b)).

**Figure 15:**
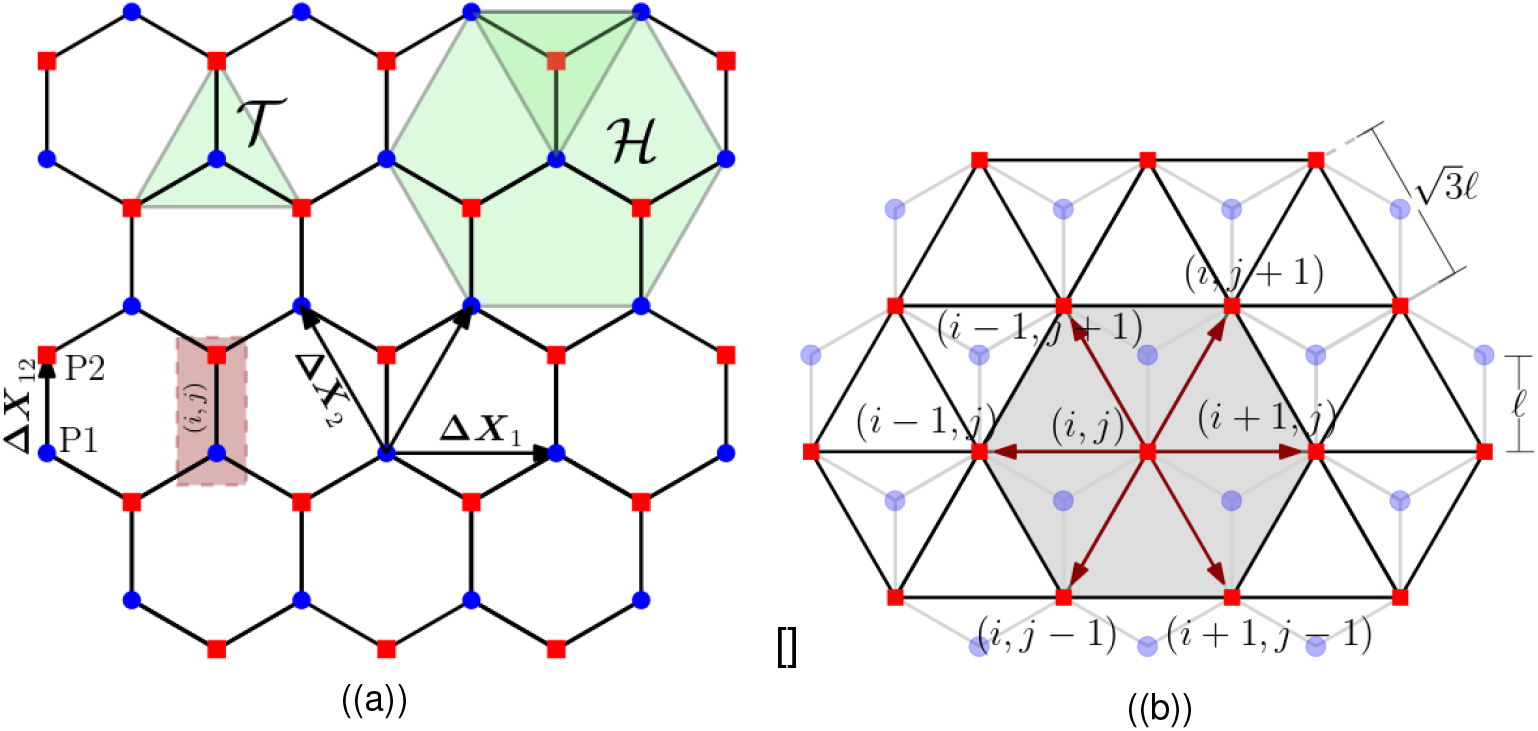
Red points (squares) are only attached to blue points (circles) and vice-versa. **(a)**: Δ***X***_1_ and Δ***X***_2_ specify the lattice directions, and Δ***X***_12_ is the edge length between Type I and Type II points. The green triangle, 𝒯 indicates the points attached to a Type I point, and the green hexagon, ℋ has as vertices the Type I points that are nearest to the Type I point at its center. **(b)**: The triangular lattice formed by subsuming blue points as secondary vertices between Type I (red square) point pairs. Dark red arrows indicate paths between nearest Type I point-pairs.

Given the lattice topology, we can now obtain a macroscale limit equation. In the asymptotic limit of many jumps, a particle has generally traveled a distance many times greater than *ℓ*, and thus we consider *ℓ* as a small parameter for an asymptotic analysis. Since the discrete Laplacian involves second-order differences, a Taylor expansion in *ℓ* near *ℓ* = 0 reduces at leading order to the Laplace-operator

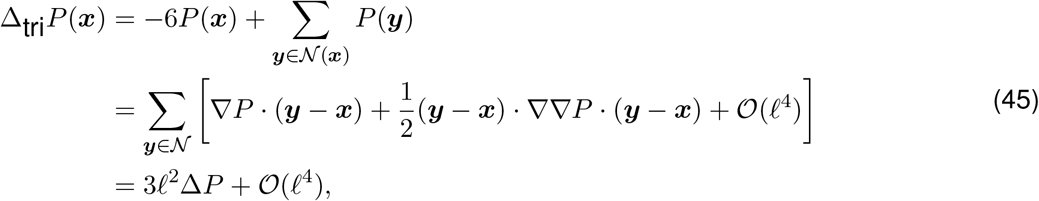

where 𝒩 (***x***) is the set of points neighboring ***x*** in a triangular lattice. All odd-order derivative terms cancel, and the factor of *ℓ*^2^ arises since all such points are a distance *ℓ* from ***x***.

We now consider the time-derivatives on the left-hand side. In the limit of many jumps occuring, we scale *ν* as *lambda* ∼ *D/ℓ*^2^ (*cf*. (Stotsky *et al*. 2021)) to obtain

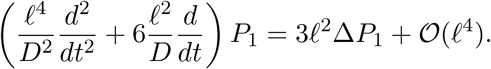

At leading order in *ℓ*, the *ℓ*^4^ terms drop out and we obtain a diffusion equation

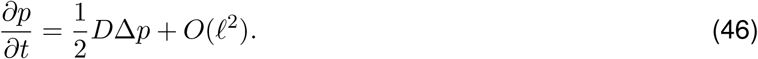

While this result is still formal since we have not proven that the higher order terms vanish, an eigenvalue analysis can be used to rigorously show that the eigenvalues corresponding to fast modes (associated with differences in the concentrations of the Type I and Type II points) vanish as *ℓ* → 0. Since there is only a single time-scale the initial ‘ballistic’ regime characteristic of the telegraph’s equation disappears as *ℓ* → 0. The fact that a telegrapher approximation never results is similar to the result in Othmer et al. (1988).

Finally, a similar Taylor series expansion shows that when *P*_1_ is continuous, *P*_2_ reduces to *P*_1_ in the small *ℓ* limit. Thus, *P*_2_ = *P*_1_ to leading order at the macroscale.

While we do not explore the diffusion case further, we do note that if a more complicated model were proposed for transport along each edge, a similar approach can be used, but as in the previous section, the diffusion coefficient becomes far more complicated.

#### 6.1.2 Transport via PIT cytonemes

To compare diffusive transport and transport by cytonemes on a hexagonal lattice, we derive a macroscale equation for a cytoneme transfer process of PIT type through the lattice. We will assume that, as with the diffusive transport described in the previous section, the cytonemes must traverse the fluid gaps between cells, and thus move along the hexagonal lattice. However, the movement of a tip is different in that the cytonemes extend and do not reverse direction until they reach a target and deliver their morphogen bolus. While each cytoneme travels without reversal, it can ’die’ along an edge in that we ignore it after it delivers its cargo to a cell. Upon reaching a junction, cytonemes cannot reverse, but can choose to travel along one of the two other edges that intersect that trijunction. While there is no reversal, cytonemes could potentially travel in a loop by making 5 right turns or 5 left turns in a row. To eliminate this possibility, which may be biologically irrelevant, we assume that cytonemes never travel in the -x direction. Under this condition, a typical cytoneme path is as shown in Figure 16. As mentioned earlier, movement may be guided by a signal emanating from receiver cells, which would automatically decrease the probability of loops in populations of cytonemes.

**Figure 16:**
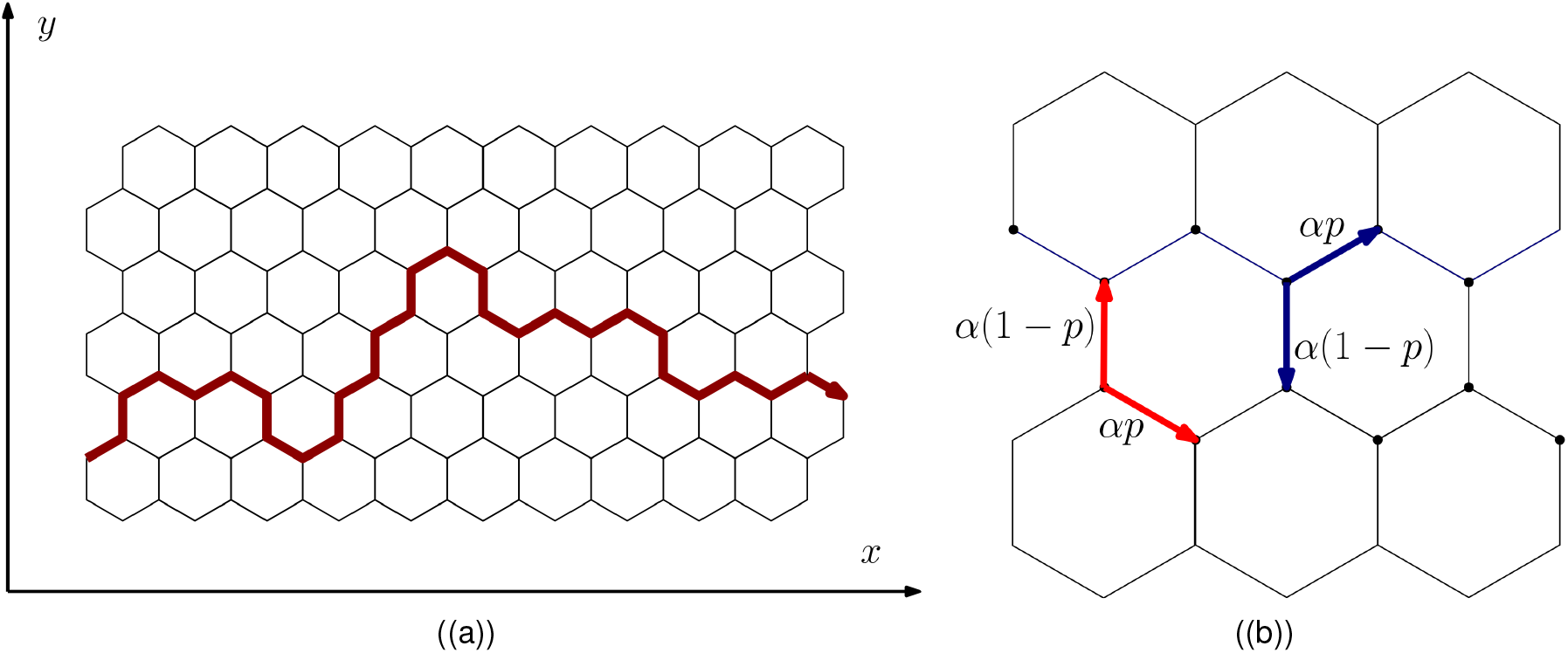
(a):The typical path of a cytoneme under the condition that it can move vertically, but always tends towards the right. (b): Spatial jump probabilities for a cytoneme departing from a Type I (blue) or fron a Type II (red) point. The two parameters, *p, α* range between 0 and 1. *p* defines the probability of jumping vertically or along an angle, and *α* gives the survival probability of the particle after each jump.

Since there are two topological types of junctions, this again leads to a 2-state system. We will assume here that the lattice is aligned as in Figure 16(a), and that all cytonemes move in the +*x*-direction except when on the vertical edges. We then have at each step a probability *αp* of continuing to the right, a probability *α*(1 − *p*) of making a vertical transition, and finally a probability (1 − *α*) for decay (see Figure 16(b)).

The evolution equations for the occupation probabilities at an arbitrary point ***x*** are

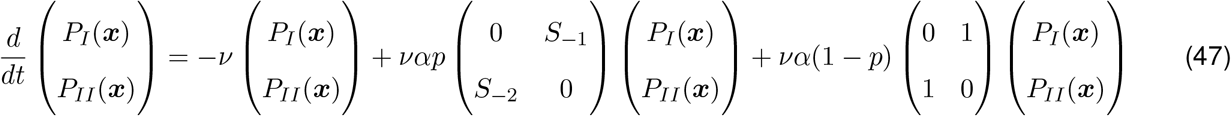

wherein [*S*_*±i*_*P* ](*x*) = *P* (*x ± δx*_*i*_) are shift operators that allow us to specify connections between adjacent points on the lattice. While it is possible to invert this 2×2 system directly, we again reduce the system to a single equation in order to determine the macroscopic equation that results from this system in the limit *ℓ* → 0. We do not expect that the distinction between Type I and Type II vertices will be significant at the macroscale, and therefore we solve for the Type II probabilities in terms of the Type I probabilities to obtain

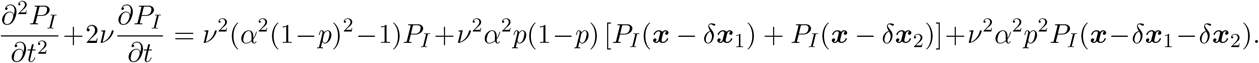

Here *ν*^−1^ is the mean time to traverse a single edge, and since the motion is advective, *ν* ∼ *v/ℓ*. We also set *α* ∼ (1 − *k*_*d*_*ℓ*) so that decay is not too fast to obtain a macroscale equation. The constant, *k*_*d*_ will be closely related to the macroscale degradation coefficient. Then, in the limit *ℓ* → 0, the leading order equation that results is

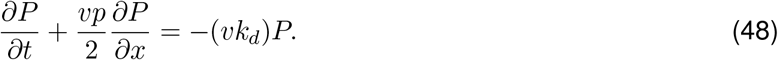

What is particularly striking about this equation is that there is no flux term in the *y* direction at large scales, despite the fact that at a microscopic scale, cytonemes can make vertical jumps. The reason is that while cytoneme makes vertical jumps up and down, on average these vertical fluctuations cancel out. Thus spreading along the *y*-axis is does not appear at leading order, and thus does not appear in the macroscopic equation.

A comparison of equations (46) and (48) shows that in the diffusive transport case, we obtain a diffusion-reaction equation with coefficients that depend on the microscale model, whereas cytoneme-based transport gives a one-dimensional advection-reaction equation. To compare these, we consider a stripe of producer cells, and for simplicity, assume that in the diffusion case, there are no-flux boundary conditions at the top and bottom of the tissue domain. If we assume uniformity in the *y*-direction, then the 2-dimensional diffusion reduces to a one-dimensional diffusion equation with a source at *x* = 0. Under these simplifications, we can compare the diffusion and cytoneme-based models as we did in the Introduction to the paper.

In each case, we impose a flux *J* at *x* = 0. For conciseness, we also set *v* and *D* equal to the macroscale diffusion and reaction coefficients obtained above, and *ν* equal to the degradation coefficients determined above. Then, for cytoneme-mediated transport, we have *J* = *vP* (0, *t*), and for diffusion, *J* = −*DP*_*x*_|_*x*=0_. The resulting solutions to this 1-dimensional problem are then as given earlier,

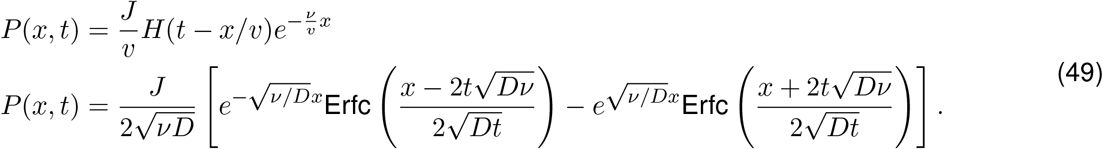

Recall that a key observation is that either can lead to exponential distributions at large times, but exhibit very different dynamics in their approach to the exponential distribution. Under diffusion, the concentration distribution gradually increases everywhere in space, whereas under advection, the concentration distribution is propagates as a travelling front, but once it is set, does not change.

When *ν* is allowed to vary, we see that for *ν <* 1, diffusion leads to a broader distribution relative to advection, whereas when *ν >* 1, the opposite occurs. Both situations have potential roles biologically, for instance, if it is important that distant cells be reached, then the strategy that yields a greater extent of travel might be more useful, whereas, if cells are detecting the gradient of a substance, then the shorter, and hence more sharply peaked distribution might be more effective at achieving biological goals.

## 7 Discussion

Herein we have developed several models of transport by cytonemes, both when the receiver cell searches for a producer cell, and when the producer cell extends cytonemes to deliver cargo to receivers. In simple models analytical solutions are attainable, and the complete spatiotemporal dependence of the morphogen concentration has been obtained. A key quantity that arises is the ratio *µ/v* of the cytoneme stopping rate to its velocity. This parameter controls the steepness of the cytoneme concentration field, and hence how far morphogen can spread.

Because it appears superficially that PIT is the most direct method of transport, one may wonder why cells employ RIT? One possibility is related to the size and shape of the target-cell domain compared with the producer-cell domain. If the latter is small, PIT could be more effective, since distant receiver cells might have difficulty finding the producer region. On the other hand, for larger or spatially-organized PIT regions, such as a boundary between tissue types, receiver cells may obtain morphogen more effectively, if, rather than waiting for a producer cytoneme to contact them, they generate their own cytonemes and thereby increase the domain which producer cells can affect. Likewise, the synaptic mode might maximize morphogen transport for cells far away from one another.

Another difference between RIT and PIT is that in the RIT-based mode studied here, the spatial distribution of morphogen approaches a constant or linearly decreasing distribution as long as all cells act similarly and have the same cytoneme production rate. This is distinct from diffusive transport and PIT, where an exponential distribution almost always results. In the PIT mode this stems from the fact that there is a finite probability that a producer cell attaches to a receiver during extension. In contrast, in RIT cytonemes always return to their origin, but if we were to introduce a finite rate of failure to return the asymptotic distribution would also be exponentially-decaying in space with a scale factor dependent on how far a cytoneme travels before it is likely to turn back.

This seems to indicate that pure RIT with no search failure is not suitable for morphogen transport since, in an undifferentiated tissue, all cells could see similar concentrations. However, this neglects the dynamics of the morphogen distribution. Cells near the producer region experience more rapid initial growth before more distant cells catch up. Thus, if the dynamics – rather than merely the amount – of morphogen is important, RIT may still be suitable. On the other hand, *perhaps cytonemes failing to find producer cells is itself a position cue that cells could use to actually measure their distance from a morphogen source*. In this sense, a hypothetical cell could generate a number of cytonemes, that perhaps even point towards multiple organizing centers, allowing a cell to in some sense triangulate its position.

In addition to RIT and PIT, we also discussed the role of puncta in morphogen transport. In essence, the repeated-packet transport mode, either as puncta within the cytoneme in PIT mode or receptor-carried packets on the surface in the RIT mode, and the single bolus mode described earlier represent two extremes of a range of transport mechanisms where continuum transport and boluses may both occur.

The simple models studied here can easily be generalized to include other effects, and exact solutions can often be obtained as long as cytonemes do not interact amongst themselves. As we have shown, stochastic simulations become appropriate for more complex models, and the models we developed, in which cytonemes, cells, and even morphogen packets are considered as separate entities, can incorporate much of what is known biologically about cytonemes. There are many further extensions that can be applied here - notably, it is straightforward to include more complicated morphogen extension and retraction steps, and the possibility of a variable morphogen bolus size. It would be of interest to ascertain experimentally whether cytonemes can interact, how the puncta sizes vary, and whether this has implications for the effects of morphogen transport.

Finally, we discussed how cell-level models that include details of the diffusive or cytoneme-mediated transport processes can be scaled up to obtain tissue-level macroscale equations. These macroscale equations take on much simpler forms, but have coefficients that depend in complex ways on the microscale details of the transport procedures. At this level, we discussed a simple comparison between diffusive transport and cytoneme-mediated transport. We found that either model can yield the same distribution of morphogen after some time, but that the dynamics of the approach to the longer-term morphogen distribution differ between the modes. This suggests that understanding the importance of diffusion and cytoneme-based transport to morphogen spreading may require not only analysis of ’steady-state’ distributions of morphogen, but also the dynamics of the morphogen concentration field over time.

## Acknowledgements

This work was supported in part by NSF Grants DMS # 185357 and # 1311974 and by NIH Grant # GM29123 to HGO. HGO would like to thank the Isaac Newton Institute for Mathematical Sciences, Cambridge, for support and hospitality during the programme Collective Behaviour where work on this paper was undertaken. This work was supported by EPSRC grant no EP/R014604/1. Any opinions, findings, and conclusions or recommendations expressed in this material are those of the authors and do not necessarily reflect the views of the National Science Foundation or of their current employers, nor do they necessarily represent the official views of the National Institutes of Health.

## 8 Appendix

### 8.1 Definition of a Poisson point-process

To define a Poisson-point process, let *N* (*t*) be the number of *events* (e.g. cytonemes generated) in [0, *t*). Each realization of the Poisson point process defines a different function *N* (*t*), or equivalently a different collection of times, 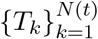 at which events occur. A Poisson process is defined by the following three properties (Ross 1996):

1. *N* (0) = 0
2. The numbers of events that occur in disjoint intervals are independent variables
3. The number of events in an interval [*s, τ* + *s*) is Poisson-distributed

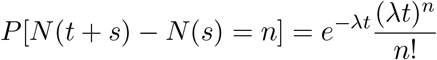

where *λ* is known as the *intensity* or *rate*. A similar result holds if *λ* varies with time.

### 8.2 Mean-values of a function of a Poisson point-process

Let *X*(*t*|*τ*_*j*_) be a function of time, *t*, and a Poisson process with arrival times *{τ*_*i*_}. Then, the following expectations can be computed in closed form,

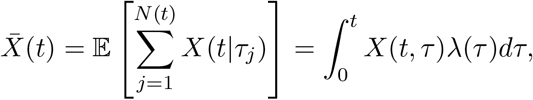

where *N* (*t*) is the number of cytonemes generated at random times *{τ*_1_, …, *τ*_*j*_} in the interval [0, *t*) and *λ*(*t*) is the Poisson-process rate parameter (which may vary in time). This result is useful in a number of cases, and can even be extended to higher order statistics using the fact that inter-arrival times are independent of one another in a Poisson process.

As an example, if cytonemes are generated at a Poisson-rate *λ*, and *q*(*x, t*) is the probability density for a single cytoneme generated at *t* = 0 to be located in (*x, x* + *dx*) at time *t*, then the expected number of cytonemes in an interval (*x, x* + *ϵ*) can be computed as

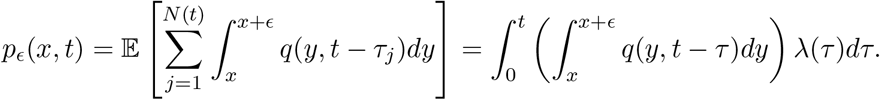

Dividing by *ϵ* and taking the limit as *ϵ* → 0, *p*_*ϵ*_ approaches

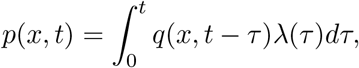

which is the number-density of cytonemes per unit length at *x*. The quantity *p*(*x, t*) satisfies the same evolution equation as *q*(*x, t*), but whereas *q*(0, *t*) = *δ*(*t*), *p*(*x, t*) satisfies the boundary condition *p*(0, *t*) = *λ*(*t*) where *λ*(*t*) is the Poisson-process rate function which characterizes the number of cytonemes generated per unit time. This is distinct from setting *p*(*x, t*) = *αe*^−*αt*^, which represents the probablity distribution for a single cytoneme. With *p*(0, *t*) = *λ*(*t*), *p*(*x, t*) is interpreted as the expected number of cytonemes per unit length at (*x, t*) instead of the probability per unit length of a single cytoneme residing in (*x, t*).

### 8.3 Cytoneme return rates and first-passage-time distributions

The fact that the expected morphogen accumulation for one or many cytonemes can be found merely by switching boundary conditions suggests an alternative interpretation to many of the results discussed thus far. In particular, the distributions for arbitrary *λ*(*t*) or *ψ*(*t*) are closely related to the solution to the equations conditioned on the cytoneme being generated at *t* = 0. In other words, given that a cytoneme is generated at *t* = 0 how long does it take for the cytoneme to reach its destination, and how much morphogen is transferred by that cytoneme? The solution to this problem is more commonly known as a first-passage-time (FPT) problem. In fact, FPT problems encompass a larger class of stochastic problems, and a classic example is to determine the distribution of times for a particle to escape from a circular region due to Brownian motion (Redner 2001)^10^. FPT’s have previously been applied to cytoneme-mediated transport in Bressloff & Kim (2019). In that context, once the FPT-distributions are known, the ‘full’ distribution for arbitrary *λ* can be found by a convolution integral. Furthermore, in subsequent sections where we consider specific times for cytoneme generation events, only the FPT is needed rather than the full distribution.

If one thinks of the FPT as the distribution of times needed to reach a given destination from a given initial point, then the FPT for the PIT process without stopping is very simple – the FPT for a cytoneme starting at *x* = 0 at *t* = 0 and stopping at *x*^*^ is simply *x*^*^*/v i*.*e*., the distribution is a delta function. If there are resting stops then the FPT is determined by the distribution of resting times. Therefore we focus on the RIT mode, and begin by defining the FPT-distribution.

Consider a single cytoneme moving according to a stochastic process. Then the FPT distribution gives the probability distribution of times needed for a cytoneme to extend, upload some morphogen molecules or morphogen-containing vesicles, and return to the receiver cell to deliver the morphogen. This distribution can be written as

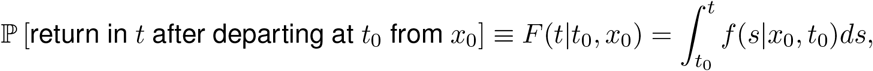

where *f* (*s*|*x*_0_, *t*_0_)*ds* is the probability density for a cytoneme that originates at (*x*_0_, *t*_0_) to return to *x*_0_ with morphogen for the first time in the interval (*s, s* + *dt*). Given *f* (*t*|*t*_0_, *x*_0_), *F* (*t*|*t*_0_, *x*_0_) represents the cumulative probability distribution that a cytoneme has returned by time *t*.

There are several ways of computing FPT distributions. In this case, we use the fact that the density function, *f* (*t*|*x*_0_, *t*_0_) is the “flux” of cytonemes returning to the receiver cell at *x*_0_ after having originated at *t*_0_. We can obtain this flux by a slight modification of equations (28) where the extension (*p*_*e*_), retraction (*p*_*r*_), and attachment (*P*_*a*_) probabilities were obtained for a cell that generates a cytoneme at a random time drawn from an exponential distribution. If instead we condition the process on the cytoneme being initiated at *t* = 0 in that example, we obtain

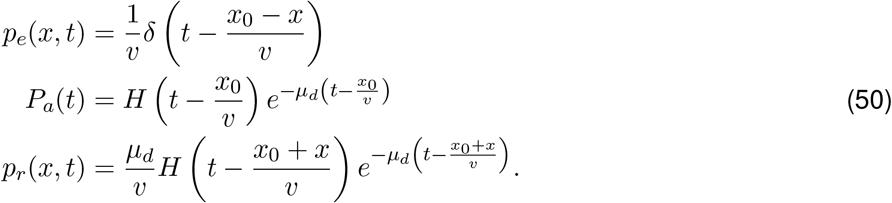

To compute the FPT distribution, we consider the fact that the probability flux for a cytoneme to reach the point *x* = *x*_0_ in the interval (*t, t* + *dt*) is equal to *vp*_*r*_(*x*_0_, *t*), and that this is equivalent in the limit as *dt* → 0 to the cytoneme experiencing a first-passage event at *t*. Thus, we have that *f* (*t*|*x*_0_, *t*_0_ = 0) = *vp*_*r*_(*x*_0_, *t*). If we now condition on *t* = *t*_0_ for some *t*_0_ ∈ R, then the result becomes *f* (*t*|*x*_0_, *t*_0_) = *vp*_*r*_(*x*_0_, *t* − *t*_0_) for any *t > t*_0_, or

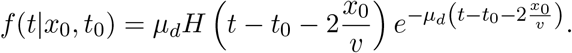

The probability of receiving morphogen is then equal to the cumulative distribution for one cytoneme, which is

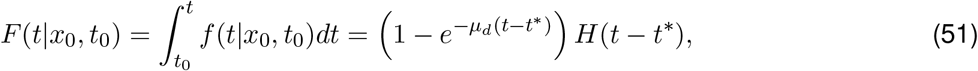

where *t*^*^ = *t*_0_ + 2(*x*_0_*/v*). A simple explanation is that because the cytonemes do not pause or reverse direction, the Dirac-delta describes the first leg of the RIT mode. This is followed by an exponentially distributed attachment time followed by return to *x*_0_. Since *F* (*t*|*x*_0_, *t*_0_) represents the accumulation of morphogen, we will denote this quantity by *m*_*F P T*_ (*t*|*x*_0_, *t*_0_) since it is analogous to *m*(*x, t*) and *m*_1_(*x, t*) used earlier. The result for *m*_*F P T*_ (*t*|*x*_0_, 0) is shown in Figure 17.

**Figure 17:**
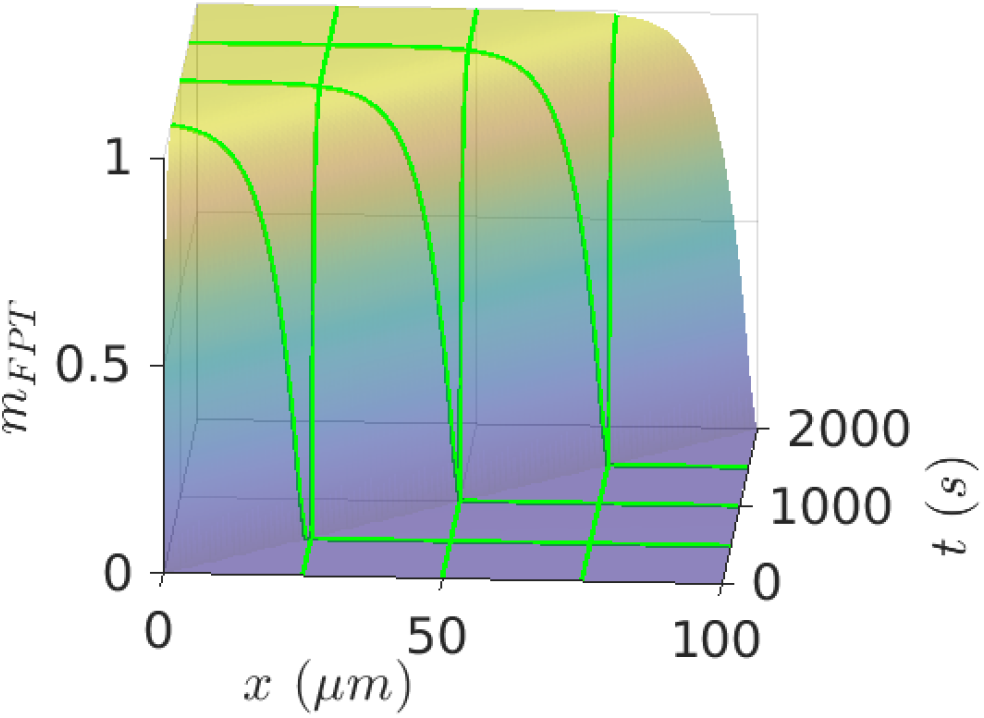
A plot of *m*(*t*|*x, t*_0_ = 0) for *λ* = 0.1*/s, µ*_*d*_ = 0.01*/s*, and *v* = 0.1*µ/s*. Moving along the *t*-axis there is a period of no accumulation followed by a rapid transition to full accumulation. Similarly, along the *x*-axis, the time delay for this to occur increases. Green lines indicate curves of constant *x* and constant *t*.

Furthermore, if we now allow for an exponentially-distributed starting time, with rate *λ*, then we recover equation (21) for the morphogen as

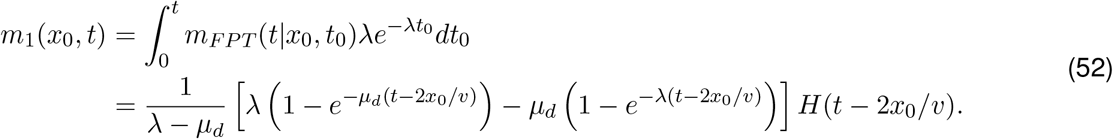

When *λ* ≪ *µ*_*d*_, the result is that cytoneme attachment is the rate-limiting factor, and when *µ*_*d*_ ≪ *λ*, cytoneme-generation is the rate-limiting factor. Likewise, *t*^*^ depends upon *x*_0_*/v*, the time it takes for a cytoneme to travel from its source to the nearest producer-cell.

We also see that in addition to *m*_1_(*x, t*), we can also recover *m*(*x, t*) from the FPT as

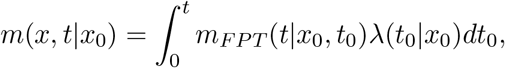

when the generation of cytonemes is governed by a Poisson process. For *λ*(*t*|*x*_0_) = *λ*_0_*H*(*t*), the resulting integral reduces to the value of *m*(*x, t*) in equation (28). This further emphasizes the importance of the FPT as a fundamental quantity since it is closely-rated to the distribution of morphogen whether we consider one cytoneme or a population.

### 8.4 The morphogen received is exponentially-distributed

Real biological systems may well exhibit deviations from the mean values computed for various cases in the main text, and it is important to understand these deviations and how cells and tissues adapt to them. The first-passage-time (FPT) distribution is useful in this regard since, under assumptions that cytonemes do not interact with each other and have uncorrelated starting times (*i*.*e*., they move and act independently), the entire probability distribution for the amount of morphogen received by each cell can be computed, rather than just its mean value. Said otherwise, in the main text we have computed

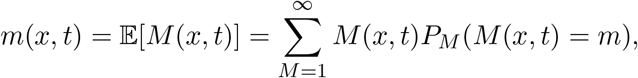

where *M* (*x, t*) is as defined in equation (7). However, it is possible to also compute the probability *P*_*M*_ (*m* = *M, x, t*), the probability for a cell at *x* to receive *M* units of morphogen packets by time *t*. Of course, in general this probability could be extremely complicated, so the assumption that cytonemes do not interfere with each other is needed if we hope to say anything analytically.

The results follow analyses described in (Ross 1996) and the previous appendices. The details of the calculation are fairly tedious, so we only present the main outcomes here. The key statistics of interest in this setting are the distributions for the number of morphogen packets and the amount of morphogen received by a cell by time *t*. When the amount of morphogen per packet is constant, these distributions are essentially the same.

If we assume that the cytonemes do not interact, these quantities are closely-related to the first-passage-time for a single cytoneme to extend, obtain a morphogen packet, and then retract. In particular, as derived earlier, the distribution of the number of morphogen packets received by a cell at *x*_0_ by time *t* is Poisson distributed with rate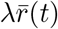:

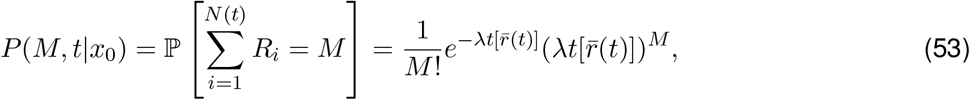

where 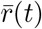 – which is discussed in detail below – is the probability for an individual cytoneme, generated at *τ* ∈ (0, *t*) to return to the originating cell with a morphogen packet by time, *t*, and *λ* is the cytoneme generation rate of the cell. If the amount of morphogen per packet is variable, then the distribution for the amount, *M* of morphogen is

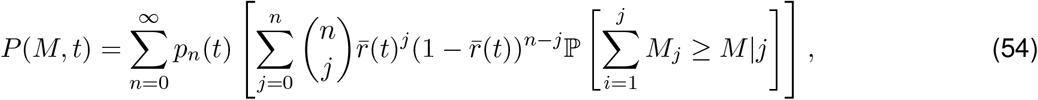

where *p*_*n*_(*t*) is the probability of generating *n*-cytonemes in (0, *t*), and the conditional probability

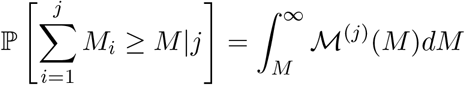

is conditioned on *j*-cytonemes returning to the cell by time *t*. The symbol ℳ^(*j*)^ is the *j*-fold convolution of the distribution function for *M* . It does not appear that this sum can be simplified to obtain a closed-form for the distribution, but can likely be approximated numerically in many cases once ℳ(*M*) is known.

One aspect of these results that is striking is that when cytonemes do not interact the morphogen-accumulation statistics depend on the cytoneme dynamics only through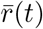, which is defined as

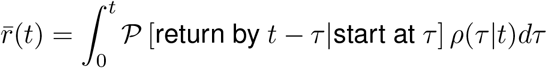

where *ρ*(*τ*|*t*) is the conditional probability for a cytoneme to be generated at *τ* given that a cytoneme is generated in [0, *t*). It is defined as

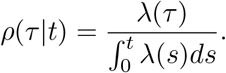

When the Poisson process rate *λ* is a constant, this reduces to *ρ*(*τ*|*t*) = 1*/t*. Likewise, the probability to return by *t* − *τ* having started at *τ* is just the cumulative distribution of the first-passage-time density, or *F* (*t* − *τ*). Hence,

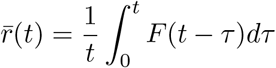

for a constant-*λ* Poisson process. The upshot is that regardless of how many different transport and binding steps an individual cytoneme undergoes, distribution of morphogen received by the receiver cells is exponential, and all of the transport steps are encapsulated in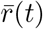.

As an example, using the cumulative FPT distribution discussed in Appendix 8.3 and computed in equation (51), we obtain

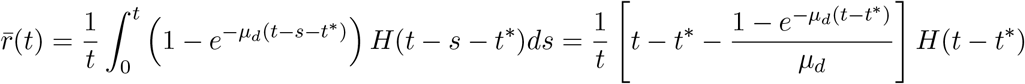

when *λ* is a constant. Note that 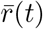 approaches 1 for large *t*, since after enough time, the cytoneme is more or less guaranteed to return. When there is a nonzero probability of failure, this will reduce the long-term limit of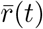.

### 8.5 Derivation of equations (53) and (54)

Here we derive the distribution of the number of morphogen packets received by a cell at *x*_0_ by time *t*, assuming that cytonemes are generated via a Poisson pointprocess of rate *λ*. The generation rate will be assumed constant, but a time-varying *λ*(*t*) can be handled via the same method of derivation with minor variations.

For a Poisson point-process, the probability of *n* events in time [0, *t*) is

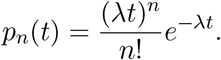

For the cytonemes, let *f* (*t*) be the FPT distribution for a cytoneme which originated at *t* = 0, to return to the cell. Then the probability for a cytoneme generated via a Poisson point-process at some time *τ* ∈ [0, *t*) to have returned to the cell is

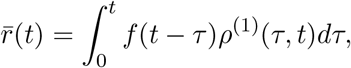

where *ρ*^(1)^(*t*) is the conditional probability of generating a cytoneme at time *τ*, given that the cytoneme is generated at some point in [0, *t*). For constant *λ*, this is just 1*/t*, and 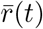 is merely the cumulative distribution of *f* (*t*) divided by *t*.

Since each cytoneme is independent, we may treat 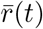 as a Bernoulli random variable, and the probability of *j* out of *n* cytonemes returning by time *t* is

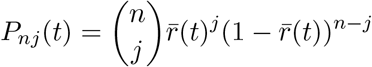

If each cytoneme carries a fixed amount of morphogen, we then write

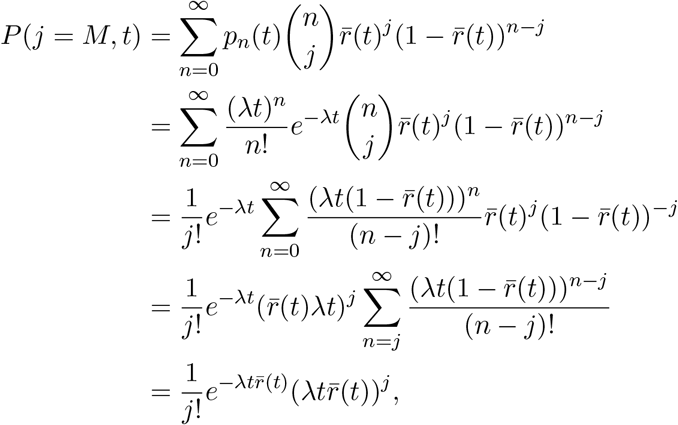

which is a Poisson distribution with rate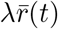. In the above derivation, we are implicitly taking all terms with *j > n* to be zero, since there cannot be more cytonemes returning then cytonemes yet generated. The derivation for variable morphogen content per packet is much the same, except that the sum now involves an addition factor that is the probability of *j*-cytonemes containing greater than *M* amount of morphogen. It does not appear that the resulting sum can be simplified as easily, though perhaps for convenient choices of the distribution of morphogen-per-packet something could be said. In this case, the probability distribution becomes,

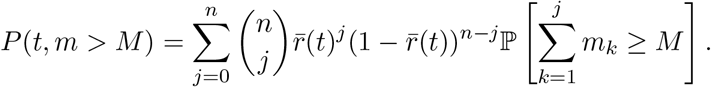

If *m* is a discrete random variable, then we can approach this by defining probabilities of the form

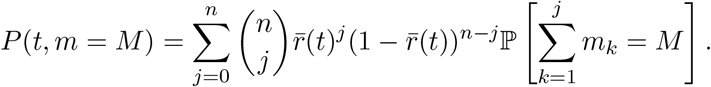

If *m* is Poisson-distributed with rate-parameter *α*, then this would lead to

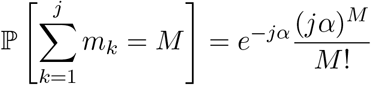

and after simplification, we obtain

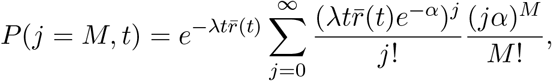

which can be computed in closed form for small *M*, and is in fact proportional to moments of a Poisson distribution. Furthermore, the moment-generating function of a Poisson distribution show that *P* (*j* = *M, t*) is related to derivatives of

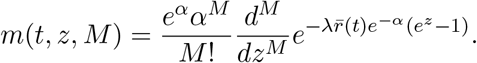

In the text, these results are applied to RIT transport, but they also turn out to apply to the PIT transport mode as well with one modification. In the PIT mode, the quantity 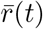 is not the probability of a cytoneme to extend and return to *x*_0_ in (0, *t*). Rather, it is the probability for a producer-generated cytoneme to extend and stop in a small interval (*x, x* + *δx*) between (*t, t* + *δt*). Since *x*(*t*) = *v*(*t* − *t*_0_) where *t*_0_ is the time cytoneme was generated, it is only necessary to consider the starting and stoping time of a cytoneme to determine when a cytoneme has stopped in (*x, x* + *dx*).

### 8.6 Simple CTRW-to-macroscale equation example

This section contains information from Example 4 of **I**. Consider a one-dimensional row of cells that are positioned so that the cell membranes are directly adjacent to one another. Let us assume that the time it takes for a molecule to transport across the membranes of the cells is Poisson-distributed with rate-parameter *λ*. Let us also assume that the time it takes the molecule to diffuse through the center of a cell is Poisson-distributed with rate *µ* (this is an approximation since the actual waiting-time distributions for diffusion are more complex, but it turns out that on a macroscale, this difference is unimportant if we set *µ* = 4*D/ℓ*^2^ where *ℓ* is the cell diameter).

This is then a classic ‘alternating’ random walk process. In particular, the following ODE system describes the probability for the particular to be at some point,

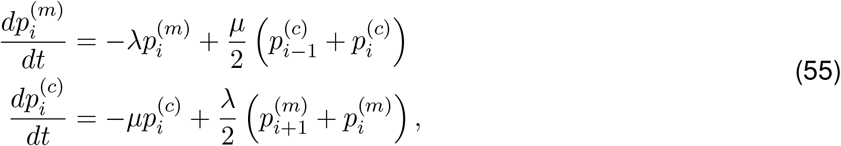

where *i* is an integer and (*m*) and (*c*) indicate membrane or cytosol of each cell.

As a first step, we consider the problem of the particle traveling from cell *i* to *i* + 1 and ignore all other cells. This is now a FPT problem for traveling from one cell to the next. It is easiest to solve this via Laplace-transform and the solution is given as

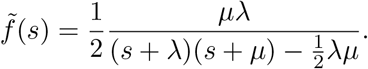

Since all cells are equivalent in this simple example, we can write a condensed CTRW with waiting-time distribution given as 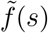 as

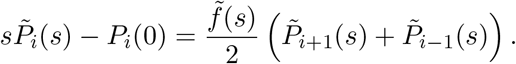

It is possible to analytically solve this problem (up to Laplace-inversion), but it is more instructive at this point to consider what the macroscale equation will be. To this end, we rearrange the equation as

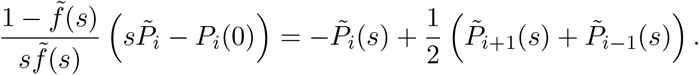

Next, we consider the limit of a cell being very small compared to the tissue, and also consider that *λ* and *µ* are rapid compared to the time-scale of interest (in other words the time and distance for a particle to move across one cell is very small). Thus, set *λ* ↦ *λ/L*^2^, *µ* ↦ *µ/L*^2^, and notice that the right-hand side of the equation is simply a discrete Laplacian, which converges to a continuum Laplacian in the small-*L* limit. Thus, we obtain to leading order

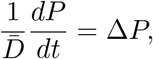

where 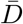 is the macroscale diffusion coefficient, defined as

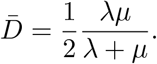

This equation now relates the microscale transport details to the macroscale behavior through the choice of 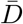. The same process can be repeated for more complicated systems, such as the hexagonal grid as well, though as the system becomes more complicated, the FPT distribution often does as well and it becomes necessary to use symbolic calculators, such as Mathematica to help with the simplification. Homogenization of a similar problem, beginning with a continuum description within each cell and the junctions, rather than a stochastic description, appears in Othmer(1983).

## Footnotes

1 A brief overview of Poisson processes and their properties relevant to the models is given in the Appendix.

2 The difficulty arises if cytonemes can interact with each other leading to effects that are not possible to capture in an individual model, or if their generation times are correlated

3 Experimental evidence suggests that sub-micron membrane fluctuations relax on a micro-to millisecond time scale (Lan & Papoian 2008) during movement, which justifies using a constant velocity.

4 One can allow the the payload to be a random variable as well, as long as it is independent of the other quantities in the model.

5 The subscript one on *m*_1_ indicates that this is for a single cytoneme.

6 The mechanism via repeated receptors illustrated in Figure (1)(a) will be treated later.

7 If there is a probability of an attached cytoneme failing to obtain a morphogen packet, we can scale the boundary condition on *p*_*e*_ by the probability of success to obtain the population of morphogen-laden cytonemes.

8 *For an exponential distribution, αe*^−*αt*^, *r*(*t*) = *α is just the rate parameter. But generally, r*(*t*) *does not simplify so conveniently*.

9 There is a distinction between the Laplacian for vertex-to-vertex jumps as here, and center-to-center jumps. In the former each vertex depends on six neighbors, whereas in the latter there are only three neighbors (Othmer & Scriven 1971; Marris *et al*. 2023)

10 A tutorial on mean FPTs and their applications in biology is given in (Polizzi *et al*. 2016)

